# RIOK2 drives glioblastoma cell proliferation by modulating MYC through the RNA-binding protein IMP3

**DOI:** 10.1101/2020.12.07.413385

**Authors:** Alexander S. Chen, Jhomar Marquez, Nathaniel H. Boyd, Se-Yeong Oh, Riley Gulbronson, Jamie A.G. Hamilton, Emily R. Legan, Renee D. Read

## Abstract

Glioblastoma (GBM) is the most common primary malignant brain tumor and is resistant to current therapies. Here, using a combination of proteomic and genetic approaches in both *Drosophila* and human GBM models, we identify novel signaling effectors that act downstream of EGFR and PI3K signaling pathways, which drive GBM. Our results indicate that RIOK2, an atypical serine/threonine kinase, forms a complex with the RNA-binding protein, IMP3 (Imp in *Drosophila*), and that the dRIOK2/RIOK-Imp/IMP3 pathway is necessary for tumorigenesis. Furthermore, our results indicate that RIOK2 catalytic activity is important for neoplasia and recruits TORC2 to phosphorylate IMP3. Finally, the dRIOK2-Imp pathway regulates protein levels of dMyc, a known Imp target mRNA, and these key findings were recapitulated in human GBM models. Collectively, our data indicates that RIOK2 catalytic activity promotes the activation of TORC2 and recruitment of IMP3, which in turn, modulates MYC mRNA and protein levels to promote tumorigenesis.

## INTRODUCTION

Glioblastoma (GBM), which is classified by the World Health Organization Classification System as grade IV neoplasms (1), is the most common adult primary malignant brain tumor. GBM originates from glia and glial progenitor cells and characteristically shows rapid cell proliferation and extensive brain infiltration. With current treatment, surgery followed by radiation and temozolomide therapy, the median survival for GBM patients remains poor at 12-15 months (2). To better understand the basic biology of GBM in order to develop improved treatments for this disease, a concerted effort has been made to genomically characterize mutations in GBM and to identify the biological mechanisms by which these mutations drive tumorigenesis.

Genomic analyses reveal that common genetic lesions in GBM alter receptor tyrosine kinase/RAS/phosphatidylinositol-3 kinase (RTK/RAS/PI3K) signaling pathways, occurring in 88% of GBMs (3). The RTK Epidermal Growth Factor Receptor (EGFR) is amplified in 40-50% of GBM cases and is typically subject to gain-of-function mutations (3, 4), the most common of which is EGFR^vIII^ (5), leading to constitutive activation of the kinase which conveys proliferative and survival advantages (6, 7). Both wild-type EGFR and EGFR^vIII^ signal through the PI3K pathway, which is commonly dysregulated in GBM (8). Genetic lesions that activate components of the PI3K pathway typically occur in 90% of GBMs (3), often as a consequence of inactivating mutations in PTEN phosphatase or activating mutations in PIK3CA and PIK3R1 (3), resulting in constitutive activation of PI3K signaling. To understand how these mutations specifically drive initiation and progression of GBM, there is a need to identify novel biological processes activated downstream of EGFR-PI3K signaling in glia. Through understanding novel mechanisms and new factors involved in GBM, we hope to uncover new potential therapeutic targets that may be used to develop new treatments for GBM patients.

To identify and investigate new downstream effectors of the EGFR-PI3K signaling pathways, we utilized a *Drosophila* GBM model (9). *Drosophila melanogaster* have several advantages as a model for neurologic cancers. *Drosophila* have a fully sequenced and annotated genome, versatile tools to allow for *in vivo* cell-type specific gene manipulation, and *Drosophila* neural and glial cell types are homologous to human neurons and glial cells (10–15). Furthermore, *Drosophila* have functional orthologs for approximately 75% of human genes including many involved in gliomagenesis (16, 17). In *Drosophila*, co-overexpression of constitutively active forms of EGFR and PI3K in glial progenitor cells produces invasive malignant neoplastic tumors that recapitulate key aspects of human GBM, including a reliance on gliomagenic core kinases like mTor and transcription factors like dMyc to drive tumorigenesis (9).

Using genetic screens in our *Drosophila* GBM models, our lab previously found that right open reading frame kinase 2 (RIOK2) is required for EGFR-PI3K-dependent glial tumorigenesis (18). RIOK2 belongs to the RIO family of proteins, which are conserved evolutionarily in organisms from *Archaea* to humans (19), comprised of at least four members (RIOK1, RIOK2, RIOK3, and RIOKB), and defined by the presence of a RIO kinase domain (19). RIO family members are considered atypical protein kinases in that the RIO kinase domain has sequence similarity to eukaryotic protein kinases (19, 20) but lacks the activation loop and substrate binding domains found in traditional eukaryotic protein kinases (19). Studies have shown that RIOK2 has functionally distinct roles in processing 18s rRNA and assembly and maturation of the 40S ribosomal subunit (21–24). In GBM tumor cells, RIOK2, which is overexpressed in EGFR amplified/mutant GBM cells, is upregulated in response to AKT activation downstream of EGFR signaling, and RIOK2 binds to and activates the mTOR complex 2 (TORC2), which in turn, phosphorylates AKT and maintains AKT activation to drive tumorigenesis (18). However, other downstream effectors of RIOK2 in tumorigenesis are unknown.

In this manuscript, we show that RIOK2 promotes GBM tumorigenesis in a novel mechanism apart from RIOK2’s known function in ribosome assembly by interacting with a complex of RNA-binding proteins, including the RBP IMP3 (IGF2BP3), which associates with RIOK2 in a manner dependent on catalytic activity. IMP3 belongs to the IGF2BP family of RNA-binding proteins, consisting of IMP1, IMP2, and IMP3, which are characterized by two N-terminal RNA-recognition motifs (RRMs) and four C-terminal hnRNP-K homology (KH) domains (25). IMP3, like other members of the IGF2BP family and other RIOK2-associated RBPs, regulates the localization, stability, and translation of cytoplasmic target mRNAs (25, 26). IMP3 is considered an oncofetal protein, in that IMP3 is normally highly expressed during embryogenesis and sparingly expressed in adult tissue, but is highly expressed in cancers (25). IMP3 expression acts as a biomarker for high-grade disease and poor prognosis in GBM and other cancers (26–28). Furthermore, studies show that IMP3 binds to and regulates mRNA of several well-established drivers of gliomagenesis including IGF2 and MYC (26, 28, 29). Our data here reveal that RIOK2 promotes GBM tumorigenesis by forming a complex with IMP3 to mediate phosphorylation of IMP3 by TORC2 to modulate MYC mRNA and protein levels.

## RESULTS

### RIOK2 catalytic activity promotes tumor cell proliferation

Because RIOK2 is thought to be a kinase and an ATPase, we sought to assess the importance of RIOK2 catalytic activity in promoting GBM tumorigenesis to determine if RIOK2 catalytic activity is required for its ability to promote tumor cell proliferation and survival. For this purpose, we tested catalytically-dead versions of RIOK2 in *Drosophila* and human GBM cells.

In our *Drosophila* GBM model in which glial-specific co-overexpression of constitutively active EGFR (*dEGFR^λ^*) and PI3K (*dp110^CAAX^*) orthologs causes glial neoplasia, genetic reduction of RIOK2 function by suppresses tumorigenesis. To determine if RIOK2 catalytic activity contributes to tumorigenesis, we used the Gal4-UAS system to overexpress catalytically-dead dRIOK2 (dRIOK2^123A,246A^) or, as a control, wild-type dRIOK2 (dRIOK2^WT^) in neoplastic glia in *repo>dEGFR^λ^; dp110^CAAX^* animals and compared their phenotypes to control *repo>dEGFR^λ^; dp110^CAAX^* animals that overexpressed a lacZ construct. dRIOK2^123A,246A^ overexpression, but not RIOK2^WT^ overexpression, significantly reduced both glial proliferation and brain volume in the GBM model (Figure 1A-F). In contrast, in normal wild-type glial cells, dRIOK2^123A,246A^ overexpression did not cause obvious or detectable defects in glial progenitor cells (*repo-Gal4*) or in neural stem cells (*wor-Gal4, dpn-Gal4*), and even whole animals (*actin-Gal4*) were grossly normal and viable with dRIOK2^123A,246A^ overexpression (Figure S1A-D, data not shown). Thus, dRIOK2^123A,246A^ can act as a dominant negative, and block tumorigenesis, while sparing normal cells. This suggests that dRIOK2 adopts a differential or neomorphic function in tumor cells, and that dRIOK^123A,246A^ overexpression specifically counteracts this function.

**Figure 1.**
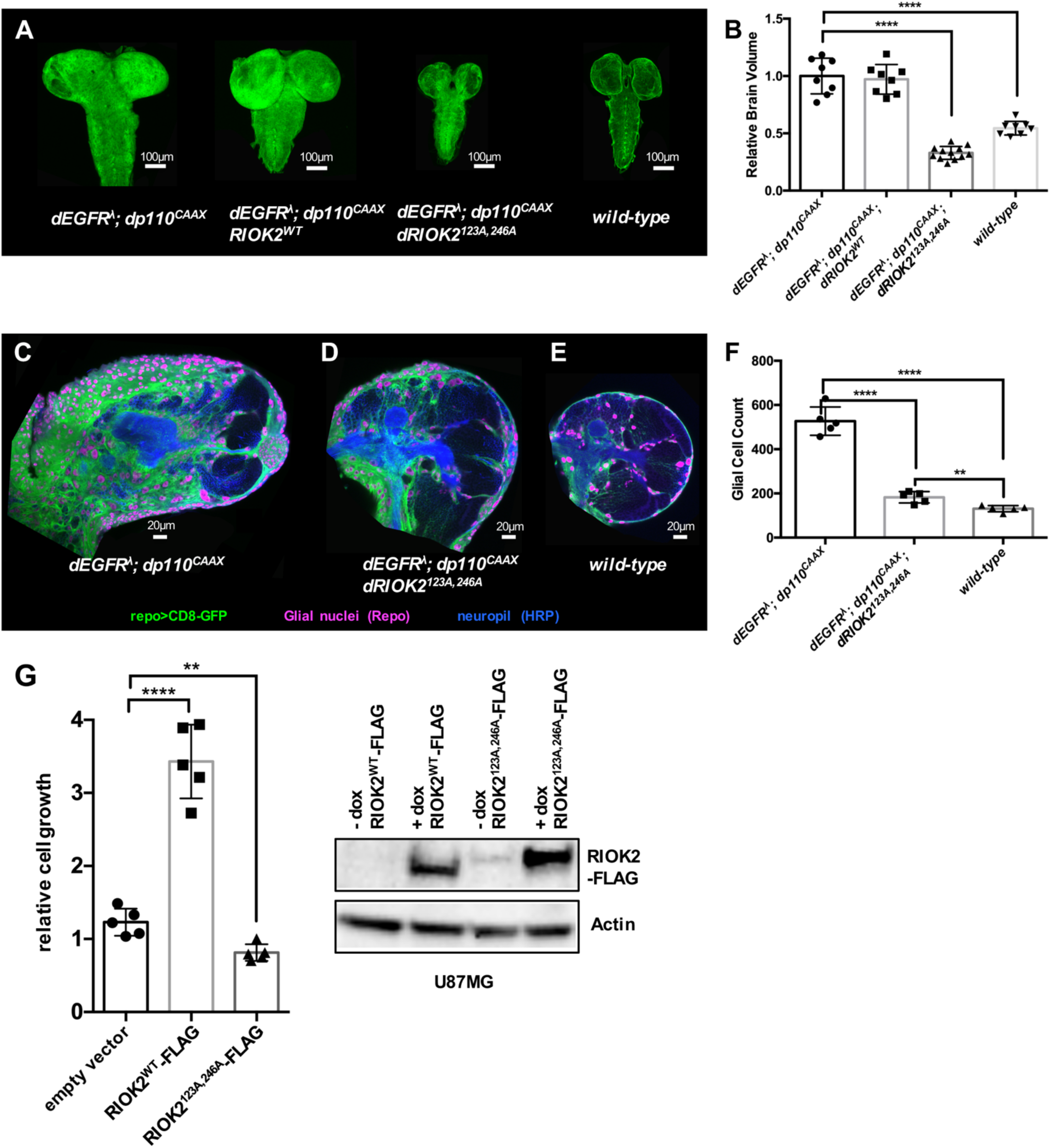
**dRIOK2/RIOK2 catalytic activity drives proliferation of tumor cells.** (A) Optical projections of whole brain-nerve cord complexes from 3^rd^ instar larvae approximately 6 days old. Dorsal view; anterior up. CD8-GFP (green) labels glial cell bodies. *repo>dEGFR^λ^;dp110^CAAX^* and overexpression of wild-type dRIOK2 (*repo>dEGFR^λ^;dp110^CAAX^*;*dRIOK2^WT^*) increased larval brain size relative to wild-type. Overexpression of catalytically inactive dRIOK2 (*repo>dEGFR^λ^;dp110^CAAX^*;*dRIOK2^123A,246A^*) decreased larval brain size relative to *repo>dEGFR^λ^;dp110^CAAX^* and *repo>dEGFR^λ^;dp110^CAAX^*;*dRIOK2^WT^* animals. (B) Total volumes of (in μm^3^) of 3^rd^ instar larval brains were measured using confocal microscopy coupled with Imaris and normalized to wild-type larval brains. (*repo>dEGFR^λ^;dp110^CAAX^*: n=8, *repo>dEGFR^λ^;dp110^CAAX^*;*dRIOK2^WT^*: n=8, *repo>dEGFR^λ^;dp110^CAAX^*;*dRIOK2^123A,246A^*: n=12, wild-type: n=8). Statistics generated using One-Way ANOVA with multiple comparisons, ****p<0.0001. (C-E) 3 μm optical projections of brain hemispheres, aged-matched 3^rd^ instar larvae. Frontal sections, midway through brains. Anterior up; midline to the left. Repo (magenta) labels glial cell nuclei; CD8-GFP (green) labels glial cell bodies; anti-HRP (blue) counterstains for neurons and neuropil. (B) *repo>dEGFR^λ^;dp110^CAAX^* and overexpression of wild-type RIOK2 (*repo>dEGFR^λ^;dp110^CAAX^*;*dRIOK2^WT^*) showed drastically increased glial cell number (magenta) and brain size compared to (E) wild-type brains. Overexpression of catalytically inactive RIOK2 (*repo>dEGFR^λ^;dp110^CAAX^*; *dRIOK2^123A,246A^*) showed drastically reduced glial cell number and decreased brain sizes compared to *repo>dEGFR^λ^;dp110^CAAX^* and *repo>dEGFR^λ^;dp110^CAAX^*;*dRIOK2^WT^* animals. (F) Glial cell numbers in representative 3 μm optical projections of 3^rd^ instar larval brain hemispheres. (*repo>dEGFR^λ^;dp110^CAAX^*: n=5, *repo>dEGFR^λ^;dp110^CAAX^*;*dRIOK2^123A,246A^* : n=5, wild-type: n=5). Statistics generated using two-tailed Student TTESTS, ****p<0.0001, **p<0.01. (G) WST-1 assays on U87MG GBM cells overexpressing either empty vector control, FLAG-tagged wild-type RIOK2 (RIOK2^WT^-FLAG), or FLAG-tagged catalytically inactive RIOK2 (RIOK2^123A,246A^-FLAG). RIOK2 constructs were induced for 48 hr by doxycycline (dox). Cultures of RIOK2^123A,246A^ -FLAG overexpressing cells displayed reduced growth compared to RIOK2^WT^-FLAG overexpressing cells, which showed increased proliferation relative to vector control cells. Five samples were used for each condition. Statistics generated using two-tailed Student TTESTS, **p<0.01, ****p<0.0001. Immunoblots confirming overexpression of RIOK2^WT^-FLAG and RIOK2^123A,246A^-FLAG upon doxycycline induction.

To examine the requirement for RIOK2 catalytic activity in human GBM cells, we tested catalytically-dead versions of RIOK2, which were created using alanine substitutions at catalytic residues Lys-123 and Asp-246 described and tested by other groups (22, 23). To determine if catalytically inactive RIOK2 (RIOK2^123A,246A^) provokes growth arrest and apoptosis in human GBM cells similar to RIOK2 RNAi-mediated knock-down, we overexpressed RIOK2^123A,246A^ in cultured cells. We had difficulty recovering GBM cells that constitutively overexpress RIOK2^123A,246A^ from viral vectors (not shown), suggesting that RIOK2^123A,246A^ blocks GBM cell growth and survival. Thus, we tested RIOK2^123A,246A^ using tetracycline inducible vectors to find that RIOK2^123A,246A^ overexpression significantly reduced growth of PTEN-null U87MG GBM cells, which are sensitive to RIOK2 levels (18), as measured by WST-1 assays (Figure 1G), whereas wild-type RIOK2 (RIOK2^WT^) overexpression enhanced cell growth, consistent with our published data showing that RIOK2 confers a proliferation advantage to PTEN mutant GBM cells (18). Thus, overexpression of catalytically-dead RIOK2 reduces proliferation of tumor cells whereas overexpression of catalytically active RIOK2 promotes proliferation. Similarly, in immortalized murine PTEN^-/-^; Ink4a/Arf^-/-^ astrocytes, constitutive overexpression of RIOK2^WT^ stimulated TORC2-Akt signaling and promoted glial tumorigenesis whereas RIOK2^123A,246A^ overexpression failed to drive TORC2-Akt signaling and did not promote tumorigenesis (18). Together, these data imply that RIOK2 catalytic function is necessary and, in the context of other oncogenic mutations such as PTEN and Inka/Arf mutation, sufficient to drive tumorigenesis.

### RIOK2 loss does not dramatically affect ribosome assembly in GBM cells

To understand how RIOK2 promotes GBM cell growth, we examined the functional impact of RIOK2 knockdown and RIOK2^123A,246A^ overexpression. In human and yeast cells, RIOK2 kinases, which are not functionally redundant with other RIO kinases, promote assembly of the 40S ribosome subunit, and loss of RIOK2 function can decrease ribosome biogenesis (21-24, 30, 31). We and others have observed that, in both *Drosophila* and human cells, RIOK2 RNAi can trigger dp53/P53 activation (18, 32). P53 transcriptional activation can occur as a result of the P53-dependent ribosome stress response, which is triggered by defects in ribosome biogenesis (33, 34). Thus, we hypothesized that, in GBM cells, RIOK2 loss or inhibition impairs ribosome biogenesis. In our cell-based GBM models, we performed sucrose gradient density analytical fractionation to characterize ribosome assembly defects caused by RIOK2 RNAi. Cells were treated with *RIOK2* shRNAi constructs combined with zVAD caspase inhibitor to block apoptosis, and lysed in hypotonic conditions so as to preserve cytoplasmic RNA macromolecules. Surprisingly, over 10 repeat experiments in three different cell types (GBM39 and GBM301 EGFR mutant GBM stem cell cultures, U87MG cell models), cells with efficient RIOK2 knockdown showed minor reductions in levels of the 40S and 60S ribosome subunit relative to monosomes as compared to control cells, and these changes were not consistent with significantly reduced ribosome assembly observed in other cell types upon RIOK2 loss (Figure S2A) (23). Of note, we observed that with RIOK2 knockdown sufficient for GBM cell growth arrest and apoptosis, residual RIOK2 protein was present in 40S subunit fractions (Figure S2B): this residual RIOK2 protein was likely sufficient for sustained ribosome biogenesis.

Our results (see above) suggest that catalytically inactive RIOK2^123A,246A^ can act as a dominant negative when overexpressed. Similar to RIOK2 RNAi, overexpression of RIOK2^123A,246A^ did not induce ribosome biogenesis defects (Figure S3A). Thus, our results indicate that reduced RIOK2 function does not primarily block tumor cell survival or growth through reduced ribosome biogenesis.

In these experiments, we examined ribosome fractions from GBM cells for other changes induced by RIOK2 knockdown. Prior studies show that TORC2 phosphorylates substrates in association with ribosomes, albeit independently of TORC2 assembly (35, 36). Because phosphorylation of TORC2 substrates is reduced by RIOK2 RNAi, we wondered if RIOK2 loss altered TORC2 association with ribosomes. We found that, upon RIOK2 knockdown, defining TORC2 components RICTOR and mTOR no longer co-fractionate with immature or mature ribosomes (Figure S2B). Thus, without substantially impacting ribosome assembly, RIOK2 knockdown impairs mTORC2 complex assembly, and thereby reduce mTORC2 activation (18). However, these data do not provide mechanistic insight into how RIOK2 modulates mTOR. While RIOK2 protein is predominantly associated with the 40S subunit peak in (Figure S2B), overexpressed cytoplasmic RIOK2 protein in GBM cells may acquire previously undescribed functions that impact TORC2.

Together with our published data showing a requirement for RIOK2 in EGFR mutant GBM cells (18), these results suggest that RIOK2 loss or inhibition affects other, as yet undiscovered, cellular processes in GBM.

### RIOK2 associates with RNA-binding proteins, including IMP3

To identify the mechanisms by which RIOK2 promotes GBM cell growth and survival, we performed immunoprecipitation (IP) experiments on both endogenous and FLAG-epitope-tagged RIOK2 protein in order to identify novel binding partners. IP experiments were carried out in RIOK2-dependent EGFR-PTEN-mutant GBM cells (U87MG-EGFR^vIII^ cells, GBM301 EGFR^vIII^ cells) under hypotonic lysis conditions to enrich for cytoplasmic RIOK2 and its binding partners. Controls were nonspecific rabbit IgG IPs and empty anti-FLAG IPs. Immunoprecipitation experiments were performed three times, and proteins associated with RIOK2 were proteomically profiled using mass spectrometry (Emory Integrated Proteomics Core) (37, 38). Datasets were compared and RIOK2 binding partners were identified that eluted with both endogenous RIOK2 and epitope-tagged RIOK2 to control for antibody-specific artifacts and eliminate non-specifically co-precipitated proteins. Potentially significant RIOK2 binding partners reproducibly met three important criteria: 1) 2-fold or greater enrichment in both endogenous RIOK2-specific IPs and FLAG-epitope-tagged RIOK2-specific IPs compared to both non-specific controls, 2) more than three spectral counts, and 3) contained 2 identified peptides.

We identified several well established RIOK2 binding proteins, including several ribosome assembly factors, including TSR1, LTV1, NOB1, and PNO1 (Figure 2A, Table S1, S2), which validates our technique (39). Furthermore, we identified several novel RIOK2 binding proteins, almost all of which were RNA-binding proteins (RBPs). The identified RBPs include IMP3, G3BP1, ATXN2L, ILF3, and others (Figure 2A, Table S1, S2), which have described primary functions in mRNA stability, transport, and translation (25). Several of these RBPs co-fractionate with RIOK2, each other, and with RBP-mRNA complexes in published studies (40, 41), increasing our confidence that they were legitimate RIOK2 partners.

**Figure 2.**
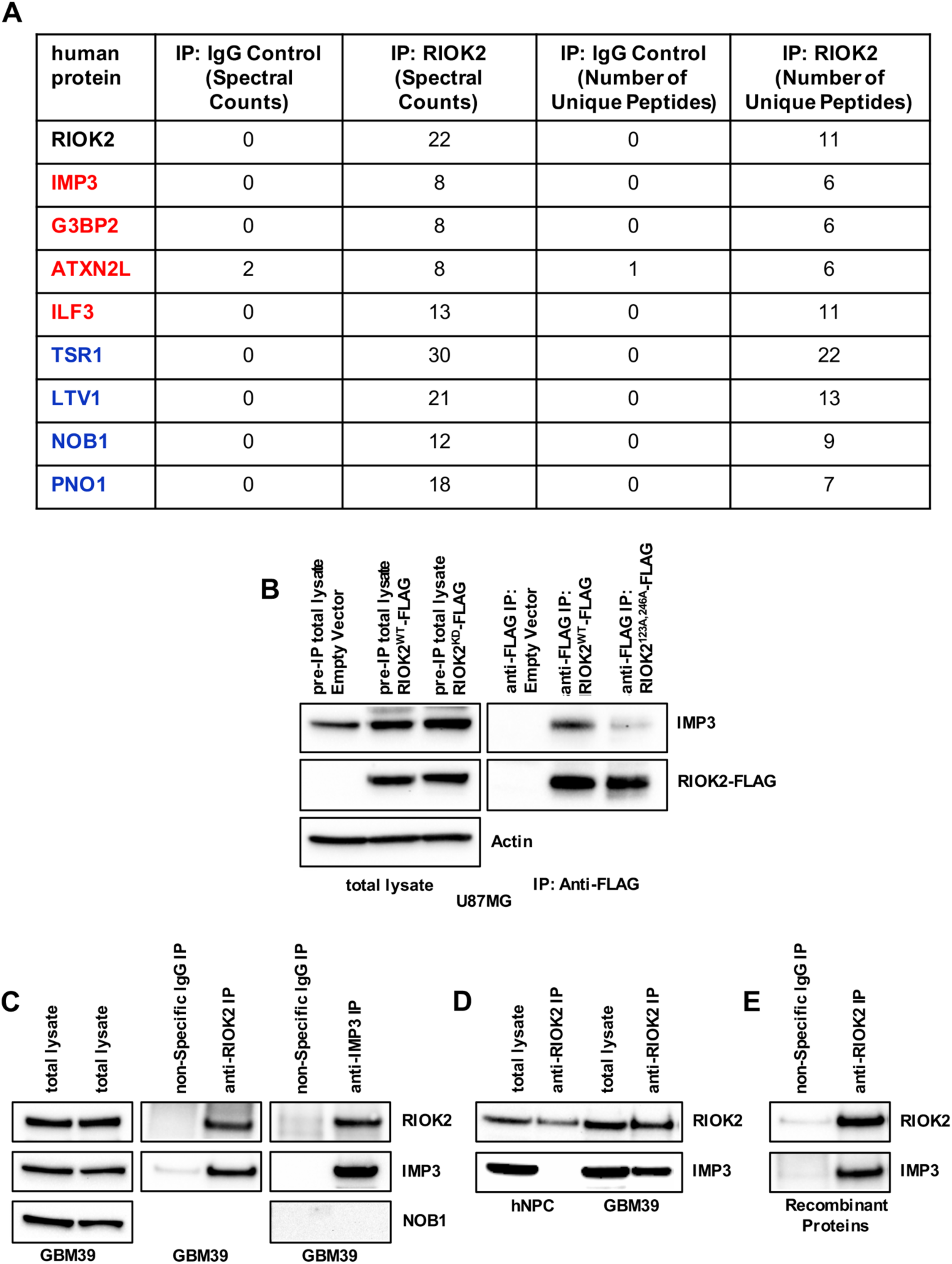
**IMP3 is a binding partner and downstream effector of RIOK2.** (A) Proteomic Analysis of RIOK2 immunoprecipitations (IPs) from U87MG-EGFR^vIII^ GBM cells performed with antibodies against either endogenous RIOK2 or with a non-specific IgG control. Values are reported in spectral counts and number of unique peptides. Spectral counts are the total number of spectra identified for a protein and is a relative measurement of the quantity of a given protein, while the number of unique peptides refers to the number of peptides that matches the protein in question. Names of RNA-binding proteins are colored red, and names of ribosome assembly factors are colored blue. (B) RIOK2^WT^-FLAG or RIOK2^123A,246A^-FLAG overexpression was induced in U87MG cells with 8 μg/ml doxycycline for 72 hrs, and proteins were IPed with Mouse M2 Anti-FLAG magnetic beads. immunoblots on total cell lysates set aside prior to IP show that endogenous IMP3 was expressed across all samples, and that FLAG-tagged RIOK2^WT^- FLAG and RIOK2^123A,246A^-FLAG were sufficiently induced by doxycycline treatment. immunoblots of show that a significantly reduced amount of IMP3 co-IPed with RIOK2^123A,246A^-FLAG compared to RIOK2^WT^-FLAG. (C) IPs from GBM39 cells using antibodies against either RIOK2 or IMP3 demonstrated reciprocal co-IP of endogenous IMP3 and RIOK2, respectively. IMP3 IPs showed no detectable co-IP of endogenous NOB1, a ribosome assembly factor and established RIOK2 binding partner. Both endogenous RIOK2 and IMP3 were detected in total cell lysates. (D) co-IPs of endogenous RIOK2 showed enrichment of endogenous IMP3 from GBM39 cells compared to human neural progenitor cells (hNPCs). Both endogenous RIOK2 and IMP3 were detected in total cell lysates of hNPCs and GBM39s. (E) Purified recombinant RIOK2 and IMP3 protein was mixed together *in vitro* and co-IPed with anti-RIOK2 antibodies, which demonstrate direct binding of IMP3 and RIOK2. RIOK2 protein and antibodies were pretreated with 0.1 mg/ml of Bovine Serum Albumin (BSA 0.1mg/ml) binding competitor prior to protein mixing and co-IP in order to reduce non-specific binding to recombinant IMP3.

To determine the contribution of RIOK2 catalytic function to RIOK2-binding partner interactions, we performed co-IP experiments on U87MG cells engineered to overexpress RIOK2^123A,246A^-FLAG compared to RIOK2^WT^-FLAG or empty vector control cells, using the same hypotonic lysis conditions described. Proteomics analysis of these co-IPs indicated that catalytically inactive RIOK2^123A,246A^ specifically co-fractionated with established RIOK2 binding partners, including ribosome assembly factors, as well as with several RBPs found to co-IP with endogenous RIOK2 and RIOK2^WT^-FLAG (Table S2). Of note, we uncovered one RBP with significantly reduced association with RIOK2^123A,246A^-FLAG relative to RIOK2^WT^-FLAG and endogenous RIOK2: the RBP IMP3 (also known as IGF2BP3) (Table S2).

To follow-up on our proteomic hits, we identified RIOK2 partners with *Drosophila* orthologs (Table S1) and then assessed the physiological relevance of these genes using a genetic modifier screen in our *Drosophila* GBM model (Table S3). The rationale for this screen being that tumor-specific functionality of RIOK2 binding partners that mediate tumorigenesis in response to EGFR-PI3K signaling should be conserved between *Drosophila* and human GBM models, and that, as with RIOK2, reduced function of these genes should modify EGFR-PI3K glial neoplasia (18). Corresponding RNAi constructs for *Drosophila* orthologs of these proteins were obtained from public sources and were tested for their ability to suppress or enhance neoplasia in our *repo>dEGFR^λ^; dp110^CAAX^ Drosophila* GBM model (18) (Table S3). For this RNAi-based modifier screen, we used a previously validated strategy in which maternally inherited Gal80 transcriptional repressor was used to reduce Gal4-mediated expression of RNAi constructs and GFP, dEGFR^λ^ and dp110^CAAX^ transgenes is delayed until later larval stages: this yield both a sensitized genetic background in the GBM model and glial-specific partial loss-of-function of RNAi-targeted genes (18). As a comparison, all RNAi constructs were also tested for their effects *in vivo* in normal *Drosophila* glia because we were particularly interested in identifying genes that were required for the growth and development of neoplastic glia but not of normal glia. We found that glial-specific RNAi of ribosome assembly factors and several RBPs significantly impaired growth and proliferation of both neoplastic and normal glia (Table S3, see Figure S4 for examples). In contrast, glial-specific RNAi of Imp, the sole IMP3 ortholog (42), significantly blocked growth and survival of neoplastic glia but had no detectable effect on the growth and survival of normal glia (see below). In this way, Imp knockdown has similar tumor-cell-specific effects as dRIOK2 knockdown and dRIOK2^123A,246A^ overexpression in *Drosophila* GBM models (see above)(18). Based on their similar phenotypic effects, we suspect that Imp and dRIOK2 may function in an evolutionarily conserved pathway in neoplastic glia.

To understand the mode of interaction between RIOK2 and IMP3 in GBM, we confirmed by immunoblot that catalytically inactive RIOK2^123A,246A^-FLAG shows reduced IMP3 binding relative to RIOK2^WT^-FLAG (Figure 2B). We also confirmed RIOK2-IMP3 binding and co-immunoprecipitation with reciprocal IPs of endogenous IMP3 with anti-IMP3 antibodies in EGFR^vIII^ mutant RIOK2-dependent patient-derived GBM stem cells (GBM39) (Figure 2C). In contrast, while control normal human neural progenitor/stem cells (hNPCs) express both IMP3 and RIOK2, but IMP3 was not associated with RIOK2 by co-IP (Figure 2D). This suggests that the interaction between RIOK2 and IMP3 is enriched in tumor cells. Of note, ribosome assembly factors such as NOB1 were not detected in the RIOK2-IMP3 IPs from GBM cells, suggesting that the RIOK2-IMP3 complex is distinct from the RIOK2-ribosome assembly factor complexes (Figure 2C). Finally, to determine if RIOK2 and IMP3 directly bind each other, we used highly validated antibodies to endogenous IMP3 and RIOK2 to perform *in vitro* co-IPs with commercially available recombinant RIOK2 and IMP3 proteins purified from bacteria: we observed co-IP of both purified IMP3 and RIOK2 proteins with each other *in vitro*, indicating that RIOK2 and IMP3 form a direct interaction (Figure 2E).

Collectively, our data indicate that RIOK2 and IMP3 are bona fide direct binding partners that may act together through a novel mechanism, perhaps with other RBPs associated with RIOK2, to drive proliferation of GBM cells, and that this mechanism may act outside of well-established RIOK2 function in ribosome assembly.

### Imp is overexpressed in EGFR-PI3K mutant neoplastic glia, and is specifically required for glial neoplasia

Imp, the *Drosophila* ortholog of IMP3, is the sole member of the IGF2BP (IMP) family of RNA-binding proteins, which is composed of three members in humans (IMP1, IMP2, and IMP3). Because of the lack of putative redundancies in *Drosophila* compared to human GBM models, to investigate the impact of knockdown of IMP family proteins in glial neoplasia we first utilized our *Drosophila* GBM model. Using immunohistochemistry, we found that Imp is upregulated in neoplastic *dEGFR^λ^; dp110^CAAX^* mutant glia compared to wild-type glia, which is consistent with previously published reports showing that IMP3 is upregulated in GBM (Figure 3A) (28, 43). We next used confocal imaging to characterize the effect of Imp functional reduction on normal and neoplastic glia. We observed that glial-specific Imp knockdown by RNAi significantly reduced both brain volume and neoplastic glial proliferation in the GBM model (Figure 3B-H). The reduction of neoplastic glia proliferation and brain volume was similar between Imp knockdown and dRIOK2 knockdown in *repo>dEGFR^λ^; dp110^CAAX^* animals, indicating that Imp is required for glial neoplasia, and could be a downstream or parallel effector of dRIOK2 (Figure 3B-H). Furthermore, Imp knockdown in wild-type glia yielded grossly normal brain morphology and glial cell development (Figure S5A-D), and animals with glial-specific Imp knockdown were viable. This result indicates that the reduction in neoplastic glial cell growth was not due to non-specific cellular lethality as a consequence of Imp knockdown, and that Imp is not essential for normal glial proliferation and development. Thus, our data suggests that similar to dRIOK2 (18), Imp is essential for proliferation of neoplastic tumorous EGFR-PI3K mutant glia, which is consistent with reports that IMP3 is required for proliferation of GBM cell lines (28, 43, 44).

**Figure 3.**
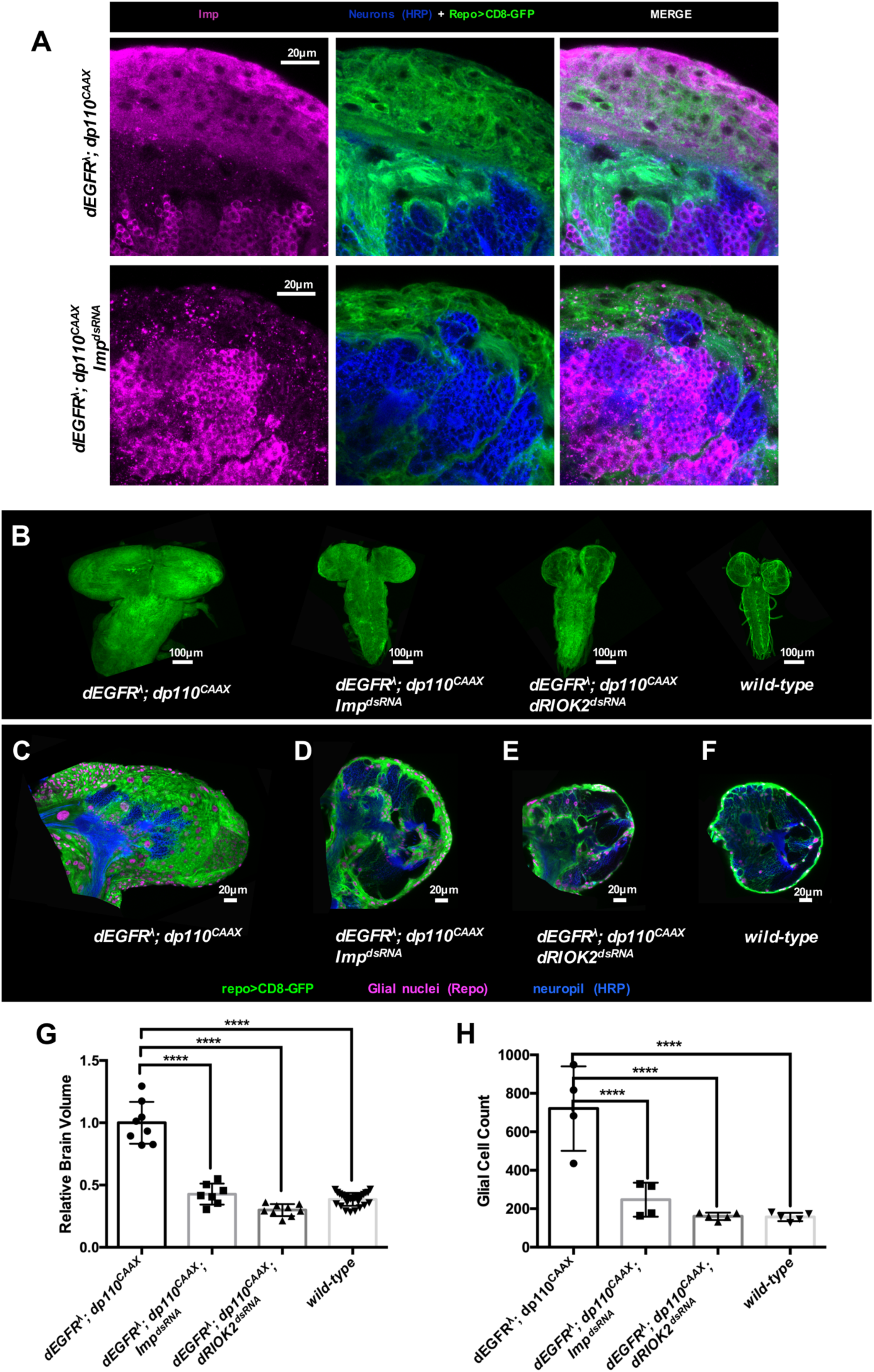
**Imp is required for glial neoplasia in a *Drosophila* GBM model.** (A) 3 μm optical projections of brain hemispheres, aged-matched 3^rd^ instar larvae. CD8-GFP (green) labels glial cell bodies; anti-HRP (blue) counterstains for neurons and neuropil; and magenta labels Imp protein (alone leftmost panels, GFP-HRP merge center panels, and Imp-GFP-HRP merge rightmost panels). Imp knockdown (*repo>dEGFR^λ^;dp110^CAAX^;Imp^dsRNA^*) (bottom panels) reduced Imp protein levels in glia (green cells in bottom panel) compared to control *repo>dEGFR^λ^;dp110^CAAX^* brains (top panels). (B) Optical projections of whole brain-nerve cord complexes from 3^rd^ instar larvae approximately 6 days old. Dorsal view; anterior up. CD8-GFP (green) labels glial cell bodies. Imp knockdown (*repo>dEGFR^λ^;dp110^CAAX^;Imp^dsRNA^*) decreased neoplastic brain overgrowth relative to *repo>dEGFR^λ^;dp110^CAAX^* controls and mimics dRIOK2 knockdown (*repo>dEGFR^λ^;dp110^CAAX^;dRIOK2^dsRNA^*), restoring brain sizes similar to wild-type animals. (C-F) 3 μm optical projections of brain hemispheres, aged-matched 3^rd^ instar larvae. Frontal sections, midway through brains. Anterior up; midline to the left. Repo (magenta) labels glial cell nuclei; CD8-GFP (green) labels glial cell bodies; anti-HRP (blue) counterstains for neurons and neuropil. (C) *repo>dEGFR^λ^;dp110^CAAX^* showed drastically increased brain sizes and glial cell number (magenta) compared to (F) wild-type brains. (C-E) Imp knockdown (*repo>dEGFR^λ^;dp110^CAAX^;Imp^dsRNA^*) and dRIOK2 (*repo>dEGFR^λ^;dp110^CAAX^;dRIOK2^dsRNA^*) significantly decreased numbers of neoplastic glia compared to *repo>dEGFR^λ^;dp110^CAAX^*. (G) Total volumes of (in μm^3^) of 3^rd^ instar larval brains were measured using confocal microcopy couples with Imaris and normalized to *repo>dEGFR^λ^;dp110^CAAX^* larval brains. (*repo>dEGFR^λ^;dp110^CAAX^*: n=8, *repo>dEGFR^λ^;dp110^CAAX^;Imp^dsRNA^*: n=7, *repo>dEGFR^λ^;dp110^CAAX^;dRIOK2^dsRNA^*: n=9, wild-type: n=27). Statistics generated using One-Way ANOVA, ****p<0.0001. (H) Glial cell numbers counted in representative 3 μm optical projections of 3^rd^ instar larval brain hemispheres. (*repo>dEGFR^λ^;dp110^CAAX^*: n=4, *repo>dEGFR^λ^;dp110^CAAX^;Imp^dsRNA^*: n=4, *repo>dEGFR^λ^;dp110^CAAX^;dRIOK2^dsRNA^*: n=5, wild-type: n=5). Statistics generated using two-tailed TTESTs, ****p<0.0001.

### RIOK2 and IMP3 are co-overexpressed and co-localized in GBM cells

Our results showed that IMP3 and RIOK2 bind to each other in a GBM tumor-cell specific manner in a manner dependent on RIOK2 catalytic activity, and that the *Drosophila* ortholog Imp is physiologically required for EGFR-PI3K-driven neoplasia in *Drosophila*. IMP3 is a known biomarker for poor patient prognosis in GBM and other cancers (26–28), although previous studies have not linked RIOK2 with regulation of IMP3 or IMP family RBPs. Therefore, we sought to further assess the relationship between IMP3 and RIOK2 in human GBM.

We first performed immunoblot analysis of a panel of patient-derived GBM neurosphere cultures created from RTK-amplified tumors, which are enriched for GBM stem cells, and observed that IMP3 is expressed at similar levels in both GBM cells and hNPCs (Figure 4A) whereas, consistent with previously observations (18), RIOK2 is overexpressed in GBM cells relative to hNPCs. This finding is consistent with previous studies showing that IMP3 expression is highest in IDH-wild-type GBM tumor tissues and is developmentally restricted to human neural stem cells (28).

**Figure 4.**
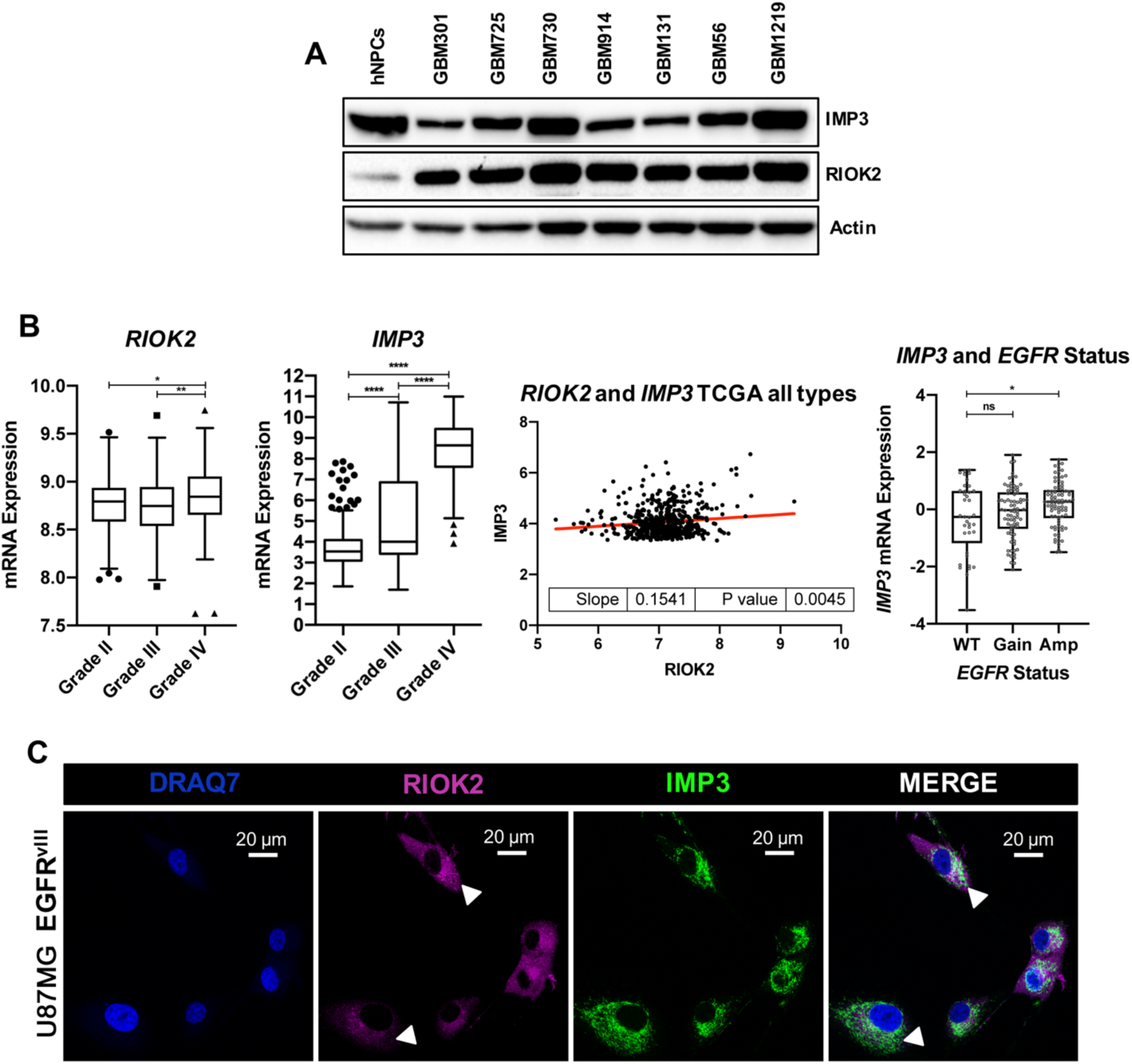
**RIOK2 and IMP3 are co-overexpressed in human GBMs.** (A) A panel of lysates from human neural progenitor cells (hNPCs) or short-term patient-derived GBM neurosphere cultures derived from tumors that showed loss of PTEN and amplification of EGFR (GBM301, GBM725, GBM914, GBM1219) or other RTKs (NTRK2: GBM131, MET: GBM730, GBM56) show elevated RIOK2 expression compared to hNPCs; all cultures express endogenous IMP3. (B) Analysis of TCGA GBM tumor profiling data show significantly increased expression of both *RIOK2* and *IMP3* mRNA in grade IV gliomas compared to lower grade gliomas. Increased expression of *RIOK2* and *IMP3* mRNA are significantly correlated with each other across all TCGA glioma grades. *IMP3* mRNA expression is significantly increased in high grade gliomas with EGFR amplification compared to tumors without EGFR amplification. (C) U87MG-EGFR^vIII^ cells were fixed and stained to label nuclear DNA (DRAQ7, blue), RIOK2 (magenta), and IMP3 (green). RIOK2 and IMP3 were co-localized predominantly in the cytosol, with both RIOK2 and IMP3 also exhibiting a perinuclear localization pattern (denoted by white arrows).

To assess whether RIOK2 and IMP3 are co-overexpressed in GBM tumor tissues, we used cBioportal and Gliovis to download and analyze tumor genomic data cataloged by The Cancer Genome Atlas (TCGA). Our data show that both *RIOK2* and *IMP3* mRNA are overexpressed to the greatest degree in grade IV gliomas (GBMs) relative to grade II and grade III gliomas (Figure 4B). Furthermore, analysis of TCGA reveal that *RIOK2* and *IMP3* mRNA expression levels are positively correlated in that high expression levels of *RIOK2* mRNA co-occur with high expression levels of *IMP3* mRNA, indicating that they may operate together to promote tumorigenesis (Figure 4B). Moreover, tumors with EGFR amplification have significantly higher expression levels of *IMP3* mRNA, indicating that IMP3 overexpression is maybe upregulated by EGFR signaling (Figure 4B). Thus, our analysis of publicly available TCGA datasets reveal that RIOK2 and IMP3 are co-overexpressed in GBM.

To investigate how RIOK2 and IMP3 regulate each other in GBM cells, we first conducted immunofluorescence experiments to assess their localization patterns in GBM cells overexpressing EGFR^vIII^. Our data show that RIOK2 and IMP3 are both predominantly localized in the cytoplasm of these cells, with both RIOK2 and IMP3 exhibiting a perinuclear staining pattern consistent with previously published IMP3 localization patterns (Figure 4C) (45, 46). These results suggest that RIOK2 and IMP3 may promote GBM tumorigenesis primarily in the cytoplasm.

Our data support that IMP3 and RIOK2 are overexpressed and co-localized in GBM tumors, consistent with the possibility that they form a functional pathway; however, the mechanism by which RIOK2 and IMP3 may work together in a pathway to promote tumorigenesis is unclear.

### RIOK2 promotes TORC2 activation and phosphorylation of IMP3

Given the requirement for RIOK2 catalytic activity for binding to IMP3 but not to the other RBPs, we first hypothesized that IMP3 is a potential substrate of RIOK2. Our *in vitro* kinase assays revealed that RIOK2 autophosphorylates and phospho-proteomic analysis identified RIOK2 autophosphorylation sites consistent with published literature (47) (Figure S6A and S6B). *In vitro* kinase assays in which RIOK2 and IMP3 were mixed followed by phospho-proteomic analysis revealed that RIOK2 does not directly phosphorylate IMP3 (Figure S6C). These results are consistent with reports that RIOK2 functions via autophosphorylation and not via direct phosphorylation of other protein substrates (23). Recent studies of RIOK2 function in 40S ribosome subunit assembly have shown that instead of directly phosphorylating substrates, RIOK2 may act more like an ATPase, wherein phosphoryl transfer from ATP to Asp257 in the RIOK2 active site and the subsequent hydrolysis of aspartylphosphate induces a conformational change that releases RIOK2 from the immature 40S ribosome subunit and from 40S assembly factors such DIM, LTV1, and NOB1 (23, 31). Therefore, we suspect that catalytically active RIOK2 may undergo a conformational change to recruit other proteins to act as intermediaries to regulate IMP3 function.

To identify additional proteins in the RIOK2-IMP3-RBP complex, we performed crosslinking IPs on endogenous RIOK2 and IMP3 in patient-derived EGFR^vIII^ mutant RIOK2-dependent GBM stem cells (GBM39) using DSP, an anime-reactive crosslinker with NHS ester reactive ends that specifically recognize proteins, combined with stringent washes, to identify transient protein-protein interactions (Figure 5A). We observed IMP3 and RIOK2 co-IP from DSP treated cells, and, by blotting for additional proteins, we found that mTOR and RICTOR, a component of mTOR Complex 2 (TORC2), were enriched in our IMP3 and RIOK2 IPs compared to non-specific IgG controls (Figure 5B). Therefore, RIOK2 may recruit TORC2 to the IMP3 protein complex.

**Figure 5.**
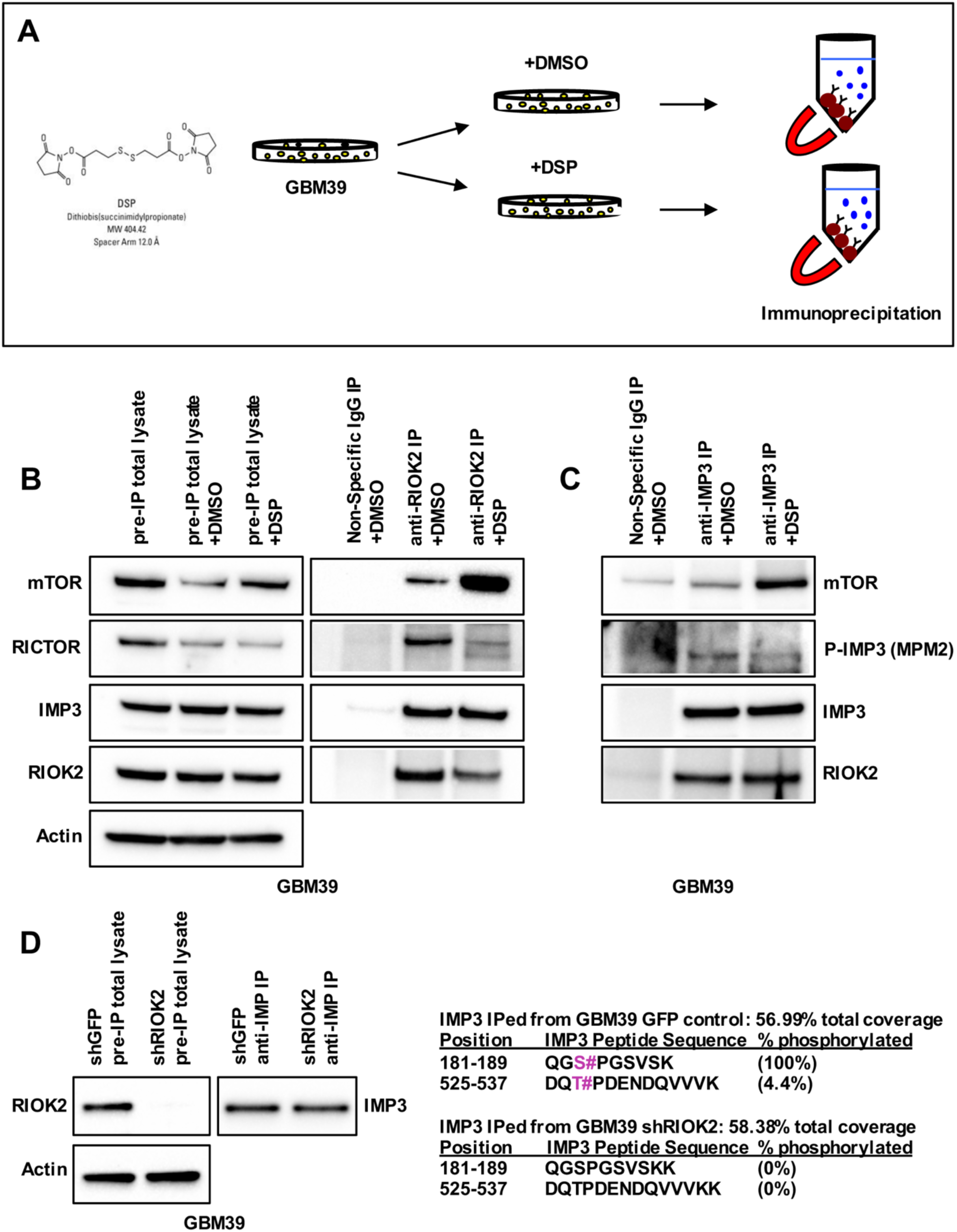
**The RIOK2-IMP3 complex recruits TORC2 and promotes phosphorylation of IMP3.** (A) Diagram of experimental workflow for DSP crosslinking and IPs from GBM39 cells. (B) GBM39 cells were treated with either DMSO or DSP crosslinker (1mM) for 2 hrs. immunoblot analysis show that GBM39 total cell lysates set aside prior to IPs endogenously expressed mTOR, RICTOR, IMP3, and RIOK2. Immunoblots of RIOK2 IPs from GBM39 cells indicate enrichment of endogenous mTOR and RICTOR protein in the RIOK2-IMP3 complex as compared to control non-specific IgG IPs. (C) GBM39 treated with either DMSO or DSP crosslinker (1 mM) for 2 hrs prior to IPs with either IMP3 antibody or a non-specific IgG control. Immunoblots show enrichment of endogenous mTOR protein in the IMP3-RIOK2 co-IPed complex, as well as phosphorylated IMP3 (detected with MPM2 antibody). (D) Phospho-peptide profiling of changes in phosphorylation of endogenous IMP3 upon RIOK2 knockdown. GBM39 cells were treated with verified lentiviral RIOK2 shRNAs. Control cells were infected with a non-targeting GFP control lentivirus. GBM cells were treated with 20 μM zVAD to prevent apoptosis. Cells were harvested 72 hrs after infection, and IMP3 was IPed. RIOK2 knockdown in pre-IP total cell lysates and anti-IMP3 IPs were verified by immunoblots. IMP3 was isolated on acrylamide gel slice and profiled for total protein and phosphorylation. Total peptide coverage for each IP in shown, and total percentage of phosphorylated peptides were calculated using according to signal intensity for phospho-peptides and total peptides for indicated fragments shown. GBM39 cells with RIOK2 knockdown showed no detectable IMP3 phosphorylation on Ser183 and Ser528 (magenta) compared to IMP3 from control cells.

Our results are consistent with our previous studies which show that RIOK2 forms a complex with TORC2, that binding of RIOK2 to mTOR depends on RIOK2 catalytic activity, and that RIOK2 catalytic activity is necessary to drive TORC2 activation (18). Previous studies show that mTOR directly phosphorylates IMP3 at Ser183 (48). To determine if mTOR phosphorylates IMP3 in the RIOK2-IMP3 complex, we performed crosslinking co-IP on endogenous IMP3 in patient-derived GBM stem cells (GBM39) followed by immunoblots with an antibody that should recognize the phosphorylated Ser183 site (MPM2, recognizes phospho-Ser/Thr adjacent to Pro), and we detected presumptive phosphorylated IMP3 in the RIOK2-IMP3-mTOR complex (Figure 5C). Furthermore, in GBM39 cells, RIOK2 knockdown reduced phosphorylation of IMP3 at Ser183 and Thr529, a second site also adjacent to a Pro residue, which supports the hypothesis that phosphorylation at these sites is regulated by RIOK2 (Figure 5D, Figure S7).

Together, our data support a model in which RIOK2 catalytic activity mediates interaction between TORC2 and IMP3, which in turn leads to mTOR-mediated phosphorylation of IMP3.

### RIOK2 and IMP3 regulate MYC levels in GBM

IMP family RBPs regulate target mRNA transport, stability, and translation (25). Previous studies in human cells show that IMP3 regulates several target mRNAs that encode established gliomagenic oncogenes including IGF2 and MYC (49). Notably, MYC is the founding member of the MYC family of “super-transcription factors,” so called because they regulate at least 15% of all known genes, many of which have established functions in a variety of pro-tumor pathways such as cell proliferation, cell growth, metabolism, ribosome biogenesis, and stem cell self-renewal (50, 51). Recent studies in *Drosophila* have identified that in neurogenesis, Imp upregulates dMyc protein levels by increasing *dMyc* mRNA stability, which in turn leads to increased proliferation and size of *Drosophila* neural stem cells, known as neuroblasts (52). Moreover, studies in human leukemia cells have shown that IMP3 binds to the 3’ UTR of *MYC* mRNA, and deletions in the RNA-binding domain of IMP3 prevented binding of IMP3 to MYC and in turn, reduced *MYC* mRNA stability (29). Together, these previous studies make a compelling case that IMP3 regulates *MYC* mRNA levels. However, no previous studies have connected RIOK2 to regulation of IMP3 or MYC.

Using our *Drosophila* GBM model, we tested whether dRIOK2 and Imp knockdown affect dMyc protein levels. Using antibodies specific to dMyc protein, our data show that dMyc protein levels are reduced with glial-specific knockdown of dRIOK2 and Imp in the GBM model compared to control animals (Figure 6A). These results are consistent with a hypothesis that a dRIOK2-Imp complex may drive glial tumorigenesis by modulating protein levels of dMyc, which is required for EGFR-PI3K driven glial tumorigenesis (9).

**Figure 6.**
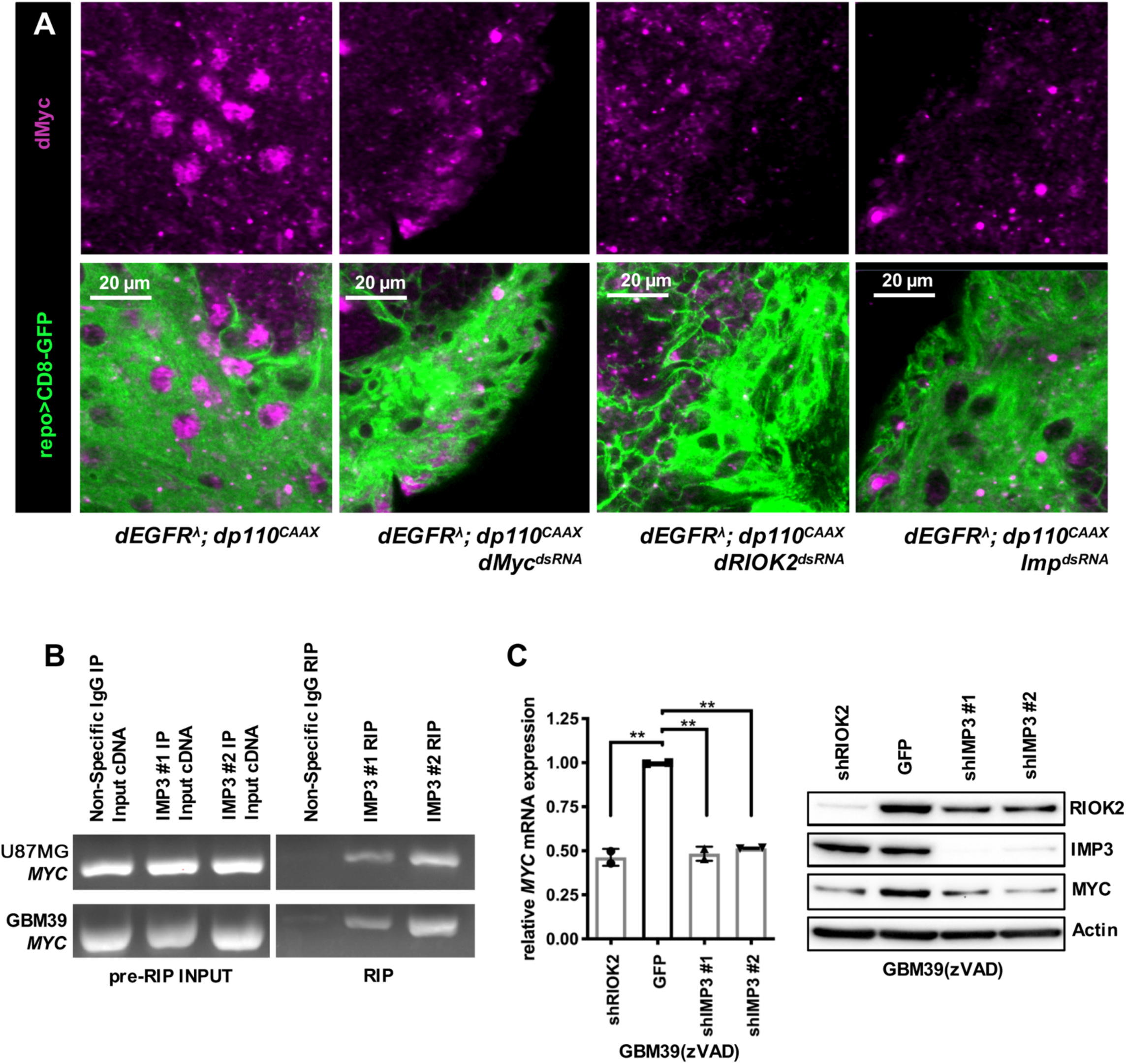
**dRIOK2/RIOK2 and Imp/IMP3 modulate dMyc/MYC expression in a conserved pathway.** (A) 3 μm optical projections of brain hemispheres, aged-matched 3^rd^ instar larvae. CD8-GFP (green) labels glial cell bodies and magenta labels dMyc protein (alone in top panels, merged in bottom panels). Glial-specific dMyc knockdown was used as a control to determine background in dMyc antibody stains. dRIOK2 knockdown (*repo>dEGFR^λ^;dp110^CAAX^*;*dRIOK2^dsRNA^*) (Third panel from the left) reduced dMyc levels in glia compared to *repo>dEGFR^λ^;dp110^CAAX^* brains (rightmost panel) and showed similar levels as dMyc knockdown(*repo>dEGFR^λ^;dp110^CAAX^*;*dMyc^dsRNA^*) (second panel of the left). Glial-specific Imp knockdown (*repo>dEGFR^λ^;dp110^CAAX^*;*Imp^dsRNA^*) (rightmost panel) reduced dMyc levels in glia compared to *repo>dEGFR^λ^;dp110^CAAX^* brains (leftmost panel) and showed similar levels as dMyc knockdown (*repo>dEGFR^λ^;dp110^CAAX^*;*dMyc^dsRNA^*) (second panel of the left). (B) IMP3 was IPed with two different antibodies (anti-IMP3 #1 and anti-IMP3 #2) from GBM39 cells. IP with non-specific IgG was used as a negative control. Co-IPed IMP3- bound mRNAs were extracted and cDNAs were generated. cDNA was PCRed with primers against *MYC*, and DNA gel electrophoresis showed specific MYC PCR products, indicating that *MYC* mRNA was bound to IMP3 in GBM cells expressing RIOK2. (C) GBM39 cells were infected with using verified lentiviral shRNAs against RIOK2 or IMP3. Control cells were infected with a non-targeting GFP control lentivirus. GBM cells were treated with 20 μM zVAD to prevent apoptosis. Cells were harvested 96 hrs after infection. GBM39 cells with RIOK2 and IMP3 knockdown showed reduced expression of endogenous *MYC* mRNA, as measured by qPCR, and MYC protein, as measured by immunoblot, compared to control cells infected with GFP shRNA. Two replicates were used per conditions. Statistics generated using unpaired two-tailed Student TTESTS, **p<0.01.

We next wanted to determine if IMP3 regulates *MYC* mRNA in human GBM cells. To determine whether IMP3 binds to *MYC* mRNA in GBM, we performed RNA-binding protein immunoprecipitations (RIP) in GBM stem cells (GBM39), using two separate IMP3 antibodies. Our data show that *MYC* mRNA is enriched in samples IPed with IMP3 compared to non-specific antibody immunoprecipitation (Figure 6B), indicating that *MYC* is a bona fide target mRNA of IMP3 in GBM cells.

To determine if RIOK2 and IMP3 regulate MYC levels in human GBM cells, we knocked down RIOK2 and IMP3 in GBM39 and GBM301 cells, and observed reduced levels of *MYC* mRNA and protein in these cells (Figure 6C, Figure S8A-B), indicating that MYC expression is upregulated by RIOK2 and IMP3. Consistent with this interpretation, analysis of TAGC data indicate that higher RIOK2 and IMP3 expression levels are significantly correlated with higher MYC mRNA expression levels, particularly in tumors that show EGFR gain and amplification (Figure S8C-D).

Collectively, our data reveal a novel mechanism in which RIOK2 recruits TORC2 and IMP3 in a complex, where mTOR phosphorylates IMP3, and together, drive GBM tumorigenesis by modulating MYC levels (Figure 7).

**Figure 7.**
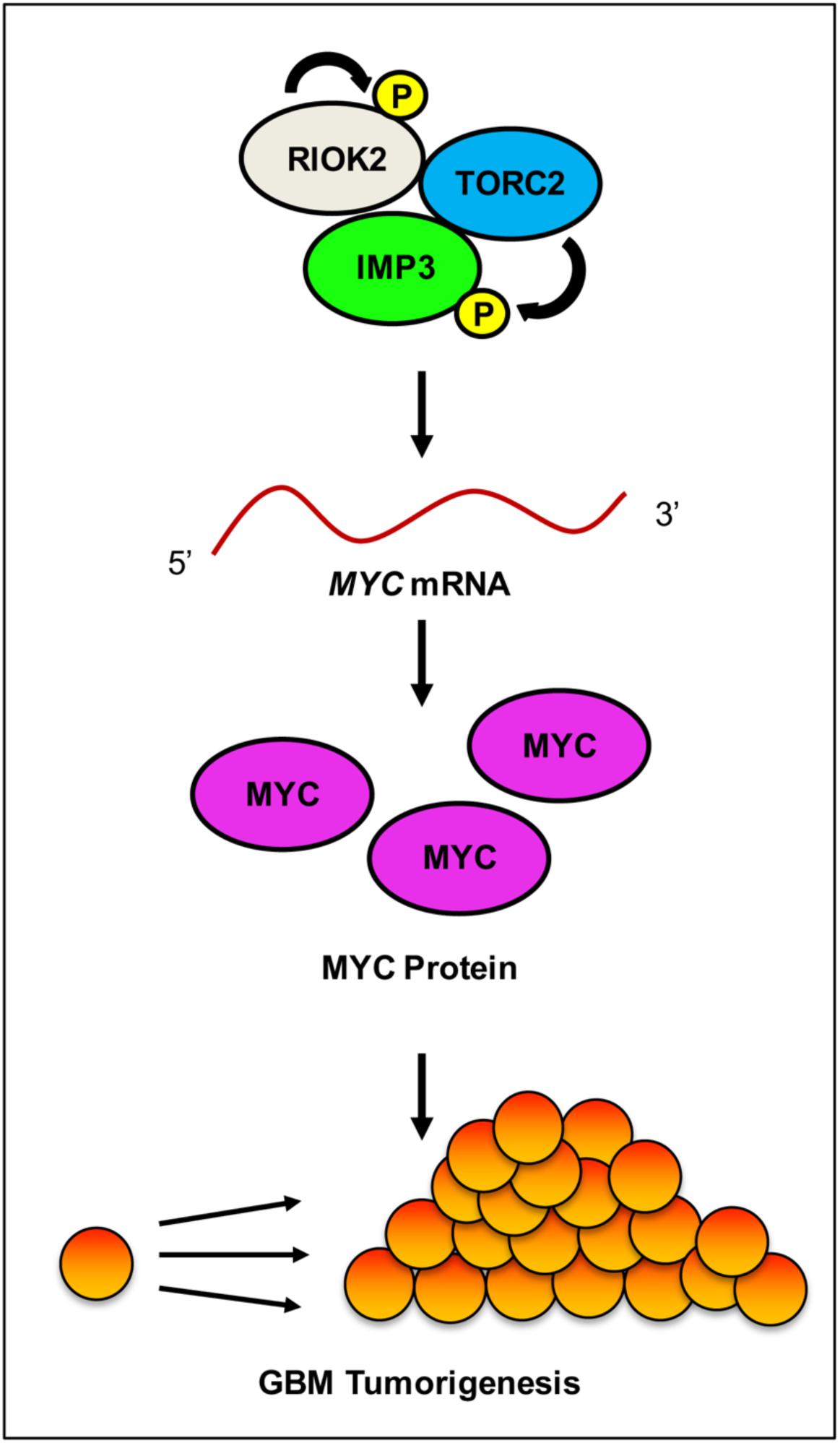
**A model for the role of the RIOK2-IMP3-TORC2 complex in regulating MYC expression to promote GBM tumorigenesis.** RIOK2 catalytic activity promotes autophosphorylation to allow for binding to IMP3 and TORC2. IMP3 is phosphorylated by TORC2 and this complex modulates *MYC* mRNA to promote increased levels of MYC protein to promote GBM tumorigenesis.

## DISCUSSION

Genes in the EGFR and PI3K pathways are commonly altered in GBM and play key roles in regulating GBM initiation and progression (53). Utilizing complementary experimental approaches in a *Drosophila* GBM model and in cell-based models of human GBM (9), we identified a new EGFR-PI3K effector pathway in which the atypical serine/threonine kinase, RIOK2, drives GBM tumorigenesis by forming a complex with the RBP IMP3, which is upregulated in tumors with EGFR amplification. The connection between RIOK2 and IMP3 is novel, and, based on what is known about IMP3, may explain the tumor cell-type-specific aspects of RIOK2 function.

Previous literature hinted at alternative functions for RIOK2 proteins aside from their conserved role in ribosome biogenesis: more than half of all Rio2p is not associated with pre-40S subunits, and, under metabolic stress, cytoplasmic Rio2p localizes to RNA granules (30, 54). Therefore, we hypothesized that RIOK2 must instead operate through a mechanism outside of ribosome assembly to promote GBM tumorigenesis. Our data supports a new model whereby RIOK2 drives GBM tumorigenesis by forming a stable and direct interaction with IMP3, which, in GBM cells, is part of a larger protein complex that includes TORC2 and likely includes other RBPs, like ATXN2, identified in our analysis of RIOK2 IPs (Table S1, Figure S9A). This concept of RIOK2 and IMP3 operating as part of larger RBP-RNA complexes to drive GBM tumorigenesis is consistent with literature supporting that RBPs act as part of messenger ribonucleoprotein particles that recruit other proteins, including kinases, to modulate their function (55, 56). In support of this, several of other RBPs that co-IPed with RIOK2 and IMP3 bind to each other and different regions on mRNAs: IMP3 and ATXN2L/ATXN2 bind to 3” UTRs and PABPC proteins bind to ATXN2L/ATXN2 and polyA tails (57, 58). Moreover, disruption of RNAs in general through RNase treatment reduced the IMP3-RIOK2 complex in cell lysates (Figure S9B), suggesting that IMP3, and likely other RBPs, coordinately bind together with RIOK2 in large-scale quaternary complexes with target RNAs.

RIOK2 has been postulated to be a kinase based on core sequence similarities between the RIO domain and other protein kinases, and has been shown to autophosphorylate itself, although no other RIOK2 substrates have been identified. Our data show that RIOK2 does not directly phosphorylate IMP3, but that RIOK2 does autophosphorylate itself, and that, in our *Drosophila* GBM model and human cell-based models, RIOK2 catalytic function is necessary for proliferation of tumorous glia. Our previous experiments reveal that RIOK2 catalytic activity is required for TORC2 binding and activation (18). Here, IP experiments in GBM cell culture show that RIOK2 catalytic activity is required for binding and formation of the RIOK2-IMP3-TORC2 complex. Therefore, we propose a model by which RIOK2 catalytic activation and autophosphorylation results in a conformational change to recruit IMP3 and TORC2, which in turn, promotes mTOR to phosphorylate IMP3, and is consistent with previous studies which have shown that, as part of the TORC2 complex, mTOR directly phosphorylates IMP3 to stimulate translation of *IGF2* mRNA (48). It is possible that RIOK2 may also promote GBM tumorigenesis through modulating TORC2 activity towards other associated RBPs, such as ATXN2L, G3BP1, G3BP2, and ILF3, and their target RNAs. Studies have shown that IMP3, along with the stress granule-associated proteins, G3BP1 and G3BP2 (59), form large aggregations of RBPs and mRNA transcripts known as stress granules (60, 61). The primary function of these granules is to store mRNA during cellular stress, such as nutrient deprivation, hypoxia, and oxidative stress (62, 63), and studies show that IMP3 provides a protective function to target mRNAs, such as *MYC,* preventing their degradation during cellular stress (61). It is possible that the elevated levels of IMP3 in GBM may protect oncogenic mRNAs from stress conditions inherent in the tumor microenvironment, further exacerbating tumorigenesis. Additional research is required to determine the mechanisms whereby RIOK2, IMP3, TORC2, and other RBPs regulate each other and larger RNA particles as well as the functional impact of those interactions on target mRNAs.

Furthermore, our *Drosophila* GBM model has identified that both dRIOK2 and the *Drosophila* ortholog of IMP3, Imp, modulates levels of dMyc, which is a key downstream effector of EGFR-PI3K signaling required for neoplastic glial transformation (9). *MYC* is a known IMP3 target mRNA (29), and MYC has an established role in gliomagenesis through upregulating expression of downstream effectors involved in ribosome biogenesis, cell proliferation, differentiation, and survival (50, 51). Importantly, our data from human GBM cells recapitulates key findings in our *Drosophila* GBM model, implicating this pathway in human disease. Our data in GBM stem cells indicate that, similarly to our *Drosophila* GBM model, RIOK2 and IMP3 co-localize, and that RIOK2 catalytic activity promotes GBM tumor cell growth and survival. Moreover, knockdown experiments in patient-derived GBM stem cells reveal that both RIOK2 and IMP3 knockdown reduced MYC mRNA and protein levels, showing that RIOK2 and IMP3 modulate MYC levels. However, the mechanisms by which the RIOK2-IMP3-TORC2 complex modulates *MYC* target mRNA are unknown. Compelling findings by Samuels *et al.* show that during development, Imp, the *Drosophila* ortholog of IMP3, stabilizes *Myc* mRNA to increase dMyc protein levels and which results in faster cell division by neuroblasts (52). In this context, RIOK2-IMP3 may operate in a similar fashion to increase MYC protein levels by stabilizing *MYC* mRNA, perhaps stimulated by TORC2- dependent phosphorylation (48). Recent studies reveal that the manner by which IMP3 regulates target mRNAs is more complicated than initially thought. Recent studies show that IMP3 regulates miRNAs in addition to mRNAs by influencing miRNA biogenesis (64), competing with miRNAs for common 3’ UTR binding sites (57), and protecting mRNAs against miRNA-dependent degradation (65). In an added level of complexity, conversely, IMP3 may promote decay of target mRNAs by promoting their association with the RISC complex, marking them for degradation (57). The ability of IMP3 to promote stability or decay of target mRNAs depends on context, such as cellular stress. Additional research is needed to elucidate the mechanisms by which the RIOK-IMP3-TORC2 complex regulates target mRNAs, such as *MYC mRNA*.

We have discovered a novel pathway by which RIOK2 promotes GBM tumorigenesis. Our data support a model wherein RIOK2 recruits IMP3 and TORC2, and TORC2 phosphorylates IMP3, and this complex promotes MYC expression to drive GBM tumorigenesis. Additional research will be needed to elucidate the mechanisms by which RIOK2 and IMP3 coordinately function with other RBPs and stress granules to modulate RNA stability, translation, and/or trafficking in GBM. This novel RIOK2 pathway may offer new avenues forward for the development of more effective therapeutics for GBM patients.

## MATERIALS AND METHODS

### *Drosophila* Strains and culture conditions

*Drosophila* stocks were obtained from the Bloomington stock center and VDRC : *UAS-Imp^dsRNA^* (Bloomington #34977, VDRC #20321, VDRC #20322); *UAS-RIOK2^dsRNA^* (VDRC #404701) ; *UAS-dMyc^dsRNA^* (VDRC #2948). *UAS-dRIOK2^WT^* and *UAS-dRIOK2* transgenic *Drosophila* were created by BestGene Inc., and multiple isogeneous stocks (>5 per construct) with independent inserts of the transgenes on all three chromosomes were bred for experimental use, tested in experimental assays, and showed nearly identical results (not shown). *Drosophila* GBM models were previously established and crosses to create tumorous larval brains were performed as described (18). All stocks were cultured on standard corn meal molasses food at 25°C. All genotypes were established by standard genetic crossing.

### Cloning

The *Drosophila* dRIOK2 MIP10156 cDNA was purchased from the *Drosophila* Genomics Resource Center and cloned into pUAS-T using standard techniques. Human RIOK2-flag cDNAs encoding isoform I (Origene) were cloned into pRev-Tre tetracycline inducible vectors (Clontech). Kinase dead catalytically inactive versions of *Drosophila* and human RIOK2 (dRIOK2^123A,246A^, RIOK2^123A,246A^) were created using site directed mutagenesis (Lightning Quick Change Kit, Agilent) to make alanine substitutions at catalytic residues Lys-123 and Asp-246 in both human RIOK2 and dRIOK2 as described by other groups (22, 23).

### Immunohistochemistry and Imaging

Larval brains were dissected with forceps, fixed in 4% paraformaldehyde, processed, stained, and imaged as previously described (9). The following antibodies were used: 8D12 mouse anti-Repo (1:10, Developmental Studies Hybridoma Bank), anti-dMyc (1:25, Developmental Studies Hybridoma Bank), or anti-Imp (1:500, a gift from Paul MacDonald, (66)). Secondary antibodies were conjugated to Cy3, Alexa-488, or Alexa-647 (1:100-250, Jackson Laboratories). Brains were mounted on glass slides ventral side down in vectashield and whole mount imaged on a Zeiss LSM 700 confocal system. For experiments where protein levels were compared between genotypes, all samples were prepared, subjected to immunohistochemistry, imaged, and image processed in a parallel manner side by side. 6 or more brains were stained with each Ab combination, and representative images are shown for each result. All brain phenotypes shown were highly penetrant, with approximately 75-100% of animals showing the growth phenotypes described. Images were analyzed in Zeiss Zen Software and processed in Photoshop. For larval brains probed with anti-dMyc, to reduce particulate background from the antibody, a median filter setting as used for the Cy3 channel with a kernel size set at 5 across all genotypes. Larval *Drosophila* brain hemisphere volumes were analyzed using Imaris software. Larval glial cells were counted manually in representative optical sections of age-matched brain hemispheres, matched for section plane. Statistical analyses were done using Prism.

### Mammalian tissue culture

GBM39, generously shared by Frank Furnari, was created from serially xenografted human GBMs. GBM725, GBM730, GBM914, GBM131, GBM56, and GBM1219 neurosphere cultures were created from GBM surgical specimens collected at Emory University and maintained in culture as described (67). Human tumor specimens were collected from surgical specimens donated for research with written informed consent of patients and were collected and used according to recognized ethical guidelines (Declaration of Helsinki, CIOMS, Belmont Report, GCP, Nuremberg Code) in a protocol (IRB00045732) approved by the Institutional Review Board at Emory University. hNPCs were obtained from Lonza. U87MG and U87MG-EGFR^vIII^ cells were gifts of Frank Furnari. Cell culture was performed as previously described (18).

Lentiviral shRNAs for RIOK2 and IMP3 (Sigma, MISSION shRNA, TRCN0000197250, TRCN0000074675, TRCN0000074675) were transfected with packaging vectors in 293 cells using Lipofectamine 3000 reagent (Thermofisher), and viral supernatant was isolated and used to infect GBM cells. For lentiviral infection, adherently plated cells were incubated with lentiviral supernatant for 6-8 hours with polybrene to enhance viral entry. Adherent neurospheres were plated in serum-free media with Geltrex (Thermofisher) or Laminin (Sigma); and adherent U87MG cells plated in 10% FBS. After infection, viral media was replaced with fresh media with zVAD caspase inhibitor if indicated, and cells were incubated for 72-96 hours prior to harvest.

U87MG cells with doxycycline-inducible wild-type RIOK2 (Rev-Tre-RIOK2^WT^) or kinase-dead RIOK2 (Rev-Tre RIOK2^123A,246A^) or empty vector (Rev-Tre alone) were created from cells engineered with the Tet3G tetracycline responsive transcriptional inducer (Clontech). Prior to collection for experiments, RIOK2 expression vectors were induced with 8 μg/ml of doxycycline and incubated for 48 or 72 hours. WST-1 assays were performed as per manufacturer’s instructions (Takara) (18).

For immunofluorescence, U87MG-EGFR^vIII^ cells were grown on glass coverslips, fixed with 4% paraformaldehyde, and stained on the coverslips with anti-IMP3 (1:100, DAKO, clone 69.1), anti-RIOK2 (1:100, Sigma HPA005681), and DRAQ7 (1:200, Cell Signaling Technology) to stain nuclear DNA and chromosomes.

### Sucrose gradient analytical fractionation for ribosome biogenesis

Ribosome profiling analysis was performed using a previously published protocol adapted for GBM cultures (68). GBM cells were cultured as described above, and to prevent apoptosis 20 μM zVAD was added for 48 hrs prior to harvest. For sucrose density gradient fractionation, hypotonic lysis (5 mM Tris pH 7.5, 2.5 mM MgCl_2_, 1.5 mM KCl, 0.1% Triton-X100 with protease and phosphatase inhibitors) was performed on 8-10 million GBM cells per experimental sample. Briefly, cells were treated with cyclohexamide and RNAseOUT (Sigma) to block translation elongation to fix ribosomes and ribosomal subunits on RNAs. Following preparation of hypotonic supernatants, the OD260 of each sample was measured (NanoDrop, ThermoFisher), and, based on OD readings, equivalent amounts of ribosomes for each lysate was loaded onto each 10%-45% sucrose density gradient (BioComp Gradient Master machine) for ultracentrifugation at 33,000 rpm for 5 hours (Beckman Optima L80K Ultracentrifuge, SW41Ti rotor). For optimal detection of ribosomes, 4-10 ODs were used per experiment, with the same ODs loaded for each control and experiments per experiment. 10% of total cell lysate was saved as an input control. Fractions were isolated and measured using a Brandel density gradient fractionator system with a UV detector, fraction collector, and chart recorder (Teledyne ISCO), and UV absorbance was recorded and analyzed with the TracerDAQ software system (MicroDAQ). Fractions were concentrated using centrifugal filters, and analyzed by immunoblot for protein composition.

### Immunoprecipitations

For immunoprecipitations used for proteomics analysis, either 2 million GBM301 cells or 5 x 10^5^ U87MG cells were lysed with hypotonic lysis buffer. 10% of the cell lysate was saved as input controls. For GBM301 and U87MG cells, 10 μg of anti-RIOK2 (1:1000, Sigma, HPA005681), or a non-specific IgG were conjugated to magnetic beads according to supplier recommendations (BIORAD, SureBeads, Protein A magnetic beads, 161-4013). In order to ensure equal amounts of protein was used for both conditions, cells were pooled and equal amounts of the pooled cells were added to each IP condition. For U87MG overexpressing RIOK2 constructs, 50 μl of anti-FLAG M2 magnetic beads (Sigma, M8823) was used. In order to ensure equal amount of protein was used per IP, cell lysates were quantified and equal amount of protein was added to each IP condition. Samples were incubated for 8 hours with end-over-end rotation at 4°C and washed five times with washing buffer (5 mM Tris pH 7.5, 2.5 mM MgCl_2_, 1.5 mM KCl). Samples were submitted to the Emory Integrated Proteomics Core for total proteomic processing according to published procedures (69). Samples for phospho-peptide mapping were submitted to the Taplin Mass Spectrometry Facility (Harvard University)

For protein-protein interactions, 2 million GBM39 cells were used per conditions and were lysed in hypotonic lysis buffer (5 mM Tris pH 7.5, 2.5 mM MgCl_2_, 1.5 mM KCl) with protease and phosphatase inhibitors as above. 10% of the cell lysis was saved as input controls. 10 μg of either anti-IMP3 (Cell Signaling 57145), anti-RIOK2 (1:1000, Sigma, HPA005681), which are highly validated by their manufacturers, or a non-specific IgG were conjugated to magnetic beads according to supplier recommendations (BIORAD, SureBeads, Protein A magnetic beads, 161-4013). In order to ensure equal amounts of protein was used for both conditions, cells were pooled and equal amounts of the pooled cells were added to each IP condition. Samples were incubated for 4 hours with end-over-end rotation at 4°C and washed five times with washing buffer (150 mM NaCl, 10 mM HEPES, 1 mM EGTA, 0.1 mM MgCl_2_, pH 7.4, and 0.1% Triton-X100). Beads were then incubated in 1X LDS sample buffer at 70°C for 10 min and analyzed using immunoblot.

*In vitro* immunoprecipitations were prepared using 0.012 μg of bacterially produced and purified recombinant RIOK2 protein (Origene, cat#TP602270) and 0.012 μg of bacterially produced and purified recombinant IMP3 protein (Origene, cat#TP760798) in hypotonic lysis buffer (5 mM Tris, 2.5 mM MgCl_2_, 1.5 mM KCl) with 0.1 mg/ml of Bovine Serum Albumin (BSA) as a competitor. 10 μg of anti-RIOK2 (1:1000, Sigma, HPA005681) or a non-specific IgG were conjugated to magnetic beads according to supplier recommendations (BIORAD, SureBeads, Protein A magnetic beads, 161-4013). Samples were incubated for 4 hours with end-over-end rotation at 4°C and washed five times with washing buffer (150 mM NaCl, 10 mM HEPES, 1 mM EGTA, 0.1 mM MgCl_2_, pH 7.4, and 0.1% Triton-X100). Beads were then incubated in 1X LDS sample buffer at 70°C for 10 min and analyzed using immunoblot.

For U87MG immunoprecipitations for tetracycline inducible systems, 5 x 10^5^ cells were plated for each condition and treated with 8 μg/ml of doxycycline for 72 hours. Cells were lysed in hypotonic lysis buffer, and 10% of total cell lysate was saved as an input control. The remaining cell lysate was quantified, and equal amounts were added to 50 μl of anti-FLAG M2 magnetic beads (Sigma, M8823) and incubated for 4 hours with end-over-end rotation at 4°C and washed five times with washing buffer (150 mM NaCl, 10 mM HEPES, 1 mM EGTA, 0.1 mM MgCl_2_, pH 7.4, and 0.1% Triton-X100). Beads were then incubated in 1X LDS sample buffer at 70°C for 10 min and analyzed using immunoblot.

### Crosslinking Immunoprecipitations

Approximately 3 million GBM39 cells were used per condition. The crosslinking immunoprecipitation was ran according to Zlatic *et al* (70). 10 μg of anti-RIOK2 (1:1000, Sigma, HPA005681), anti-IMP3 (Cell Signaling 57145), or a non-specific IgG were conjugated to magnetic beads according to supplier recommendations (BIORAD, SureBeads, Protein A magnetic beads, 161-4013). Following washes, beads were then incubated in 1X LDS sample buffer at 70°C for 10 min and analyzed using immunoblot.

### *In vitro* Kinase Assay

10 μl reactions containing 1 μg of purified recombinant protein 1 µCi γ-[^32^P] ATP in 50 mM Tris/HCl, pH 7.6, 200 mM NaCl, and 5 mM MgCl_2_ (Kinase Buffer) were incubated at 37°C for 1 hour. Samples were then immunoprecipitated for 4 hours with end-over-end rotation at 4°C using magnetic beads conjugated with anti-IMP3 (Cell Signaling 57145) according to supplier recommendations (BIORAD, SureBeads, Protein A magnetic beads, 161-4013). Samples were washed five times with kinase buffer. Beads were then incubated in 1X LDS sample buffer at 70°C for 10 min and analyzed using phosphoimager.

### Immunoblot Analysis

The following antibodies were used for immunoblotting following the manufacturer’s recommendations: anti-RIOK2 (1:1000, Sigma HPA005681; 1:1000, Origene Clone OTI3E11), anti-IMP3 (1:1000, Cell Signaling 57145; 1:1000, DAKO clone 69.1), anti-mTOR (1:1000, Cell Signaling 4517), anti-MYC (1:1000, Cell Signaling clones D3N8F and E5Q6W), anti-NOB1 (1:1000, Origene TA808793 clone OTI1C12), anti-phospho-Ser/Thr-Pro MPM-2 (1:500, Millipore-Sigma 05-368), and anti-actin (1:200, Developmental Studies Hybridoma Bank JLA20).

### *in silico* Analysis

Patient RNAseq data from The Cancer Genome Atlas (TCGA) was downloaded using Gliovis (71) (gliovis.bioinfo.cnio.es) and XY correlations were performed between *RIOK2*, *IMP3*, *MYC*, and *mTOR* mRNA expression (data from 528 GBM patients), as well as mRNA expression of *RIOK2* and *IMP3* by tumor grade (data from 620 glioma patients). Copy number analysis for *EGFR* and comparison with *IMP3* mRNA expression was performed using cBioportal (data from 201 patients).

### RNA-binding Protein Immunoprecipitation

RNA-binding protein immunoprecipitation was run according to supplier recommendations (Millipore, Magna RIP Kit, 17-700). Beads were conjugated with 10 μg either anti-IMP3 (#1: MBL, RN009P, #2: Cell Signaling, 57145) or non-specific IgG. cDNA was generated from IPed RNA, PCRed with primers against MYC, and the PCR results were analyzed using DNA gel electrophoresis.

### Statistical analyses

Larval brain volumes were analyzed using one-way ANOVA with multiple comparisons. Larval glial cell numbers were analyzed using two-tailed parametric T-Tests. WST-1 data was analyzed using two-tailed parametric T-Tests. TCGA microarray data analysis was downloaded and graphed using Graphpad Prism. Correlative data was analyzed using linear regression, and the slope and P value are reported as calculated from an F test that answers the question “is the slope significantly non-zero?” Tumor grade data and copy number analysis was analyzed using one-way ANOVA with multiple comparisons and are reported using p values. Comparisons between two groups were done using two-tailed parametric T-Tests.

## Acknowledgements

We would like to thank Coston Rowe, Colleen Mosley, Duc Duong, Avanti Gokhale, and Sara Leung for technical assistance with *Drosophila* genetics, cell culture, proteomics, protein crosslinking, and *in vitro* kinase assays, respectively, Victor Faundez and Yue Feng for intellectual guidance and helpful discussions, and Anita Corbett and Brian Petrich for critical reading of the manuscript.

## Supplementary Figures

**Figure S1.**
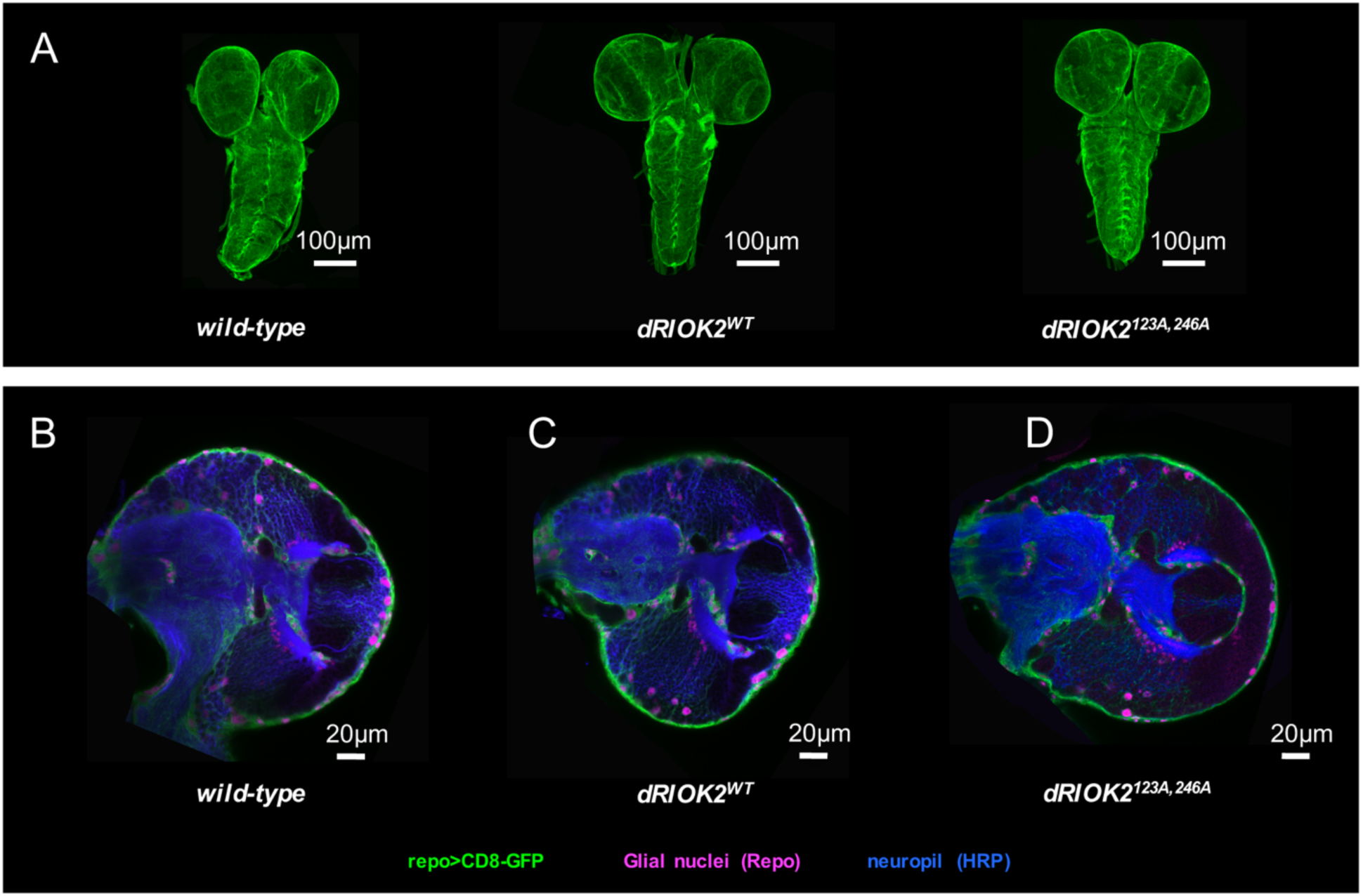
**Overexpression of wild-type dRIOK2 or catalytically inactive dRIOK2 induced no gross development or morphological defects in *Drosophila* glia.** (A) Optical projections of whole brain-nerve cord complexes from 3^rd^ instar larvae approximately 6 days old. Dorsal view; anterior up. CD8-GFP (green) labels glial cell bodies. Overexpression of either wild-type dRIOK2 (*repo>dRIOK2^WT^*) or kinase-dead dRIOK2 (*repo>dRIOK2^123A,246A^*) induced no drastic morphological differences in the brain compared to wild-type control animals. (B-D) 3 μm optical projections of brain hemispheres, aged-matched 3^rd^ instar larvae. Frontal sections, midway through brains. Anterior up; midline to the left. Repo (magenta) labels glial cell nuclei; CD8-GFP (green) labels glial cell bodies; anti-HRP (blue) counterstains for neurons and neuropil. Compared to wild-type animals (A), (B) overexpression of wild-type dRIOK2 (*repo>dRIOK2^WT^*) or overexpression of catalytically-dead dRIOK2 (*repo>dRIOK2^123A,246A^*) produced no drastic morphological differences compared to wild-type animals showed drastically different brain morphology.

**Figure S2.**
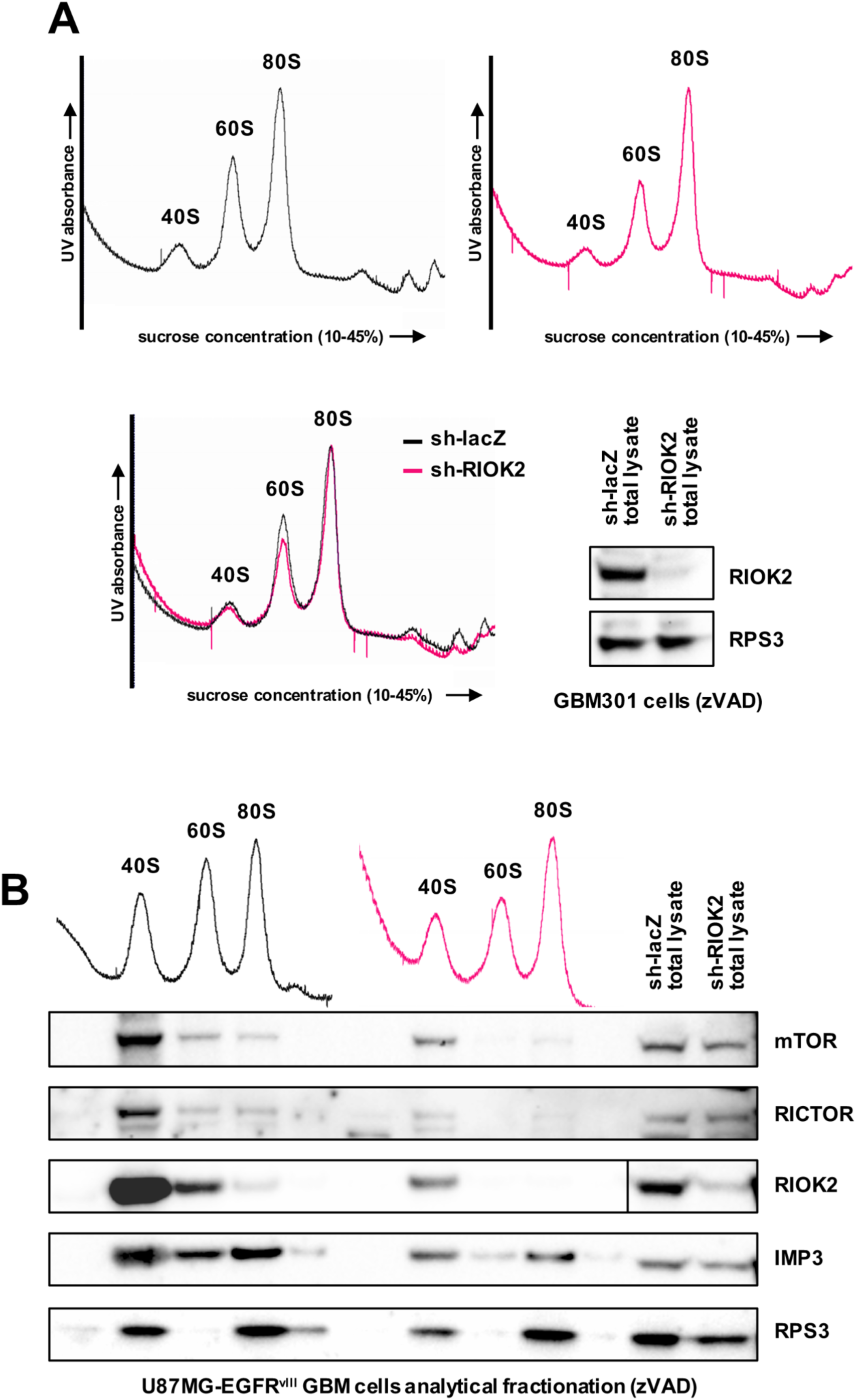
**RIOK2 RNAi induces minor defects in ribosome biogenesis and disrupts TORC2 activation.** A) Analytical sucrose density gradient fractionation of GBM301 cells treated with either a lentiviral control lacZ shRNA or a validated RIOK2 shRNA. Cells were infected with shRNAs, treated with 20 μM zVAD, and harvested 72 hrs following infection. Immunoblots of total cell lysates (right) confirmed RIOK2 knockdown and equivalent expression levels of ribosomal proteins between control and RIOK2 knockdown samples. 4 ODs from each lysate were ultracentrifuged in a sucrose density gradient of 10%-45%, and eluted into several fractions while a UV lamp was used to detect ribosome subunits. Annotated UV traces show that RIOK2 knockdown slightly reduced levels of 40S and 60S subunits relative to levels of mature 80S monosomes. B) RIOK2 co-fractionates with mTOR and RICTOR in association with the 40S ribosome subunit in GBM cells by analytical sucrose density gradient fractionation. RIOK2 knockdown efficiency confirmed by immunoblot on total cell lysate, right two lanes. Annotated UV traces for each sample (9 ODs used for each) is shown above immunoblot results for concentrated matched fractions, which show that RIOK2 knockdown induced a slight reduction in levels of 40S and 60S subunits relative to mature 80S monosomes. RIOK2, mTOR, and RICTOR proteins are enriched in the concentrated fractions collected for the 40S peak, as confirmed by the presence of the small subunit protein RPS3. RIOK2 knockdown inhibited mTOR-RICTOR co-fractionation with the 40S ribosome subunit. IMP3 also co-fractionated with mature and immature ribosomes. These results may indicate either direct association or independent co-fractionation of RIOK2-IMP3-mTOR-RICTOR complexes with ribosome subunits.

**Figure S3.**
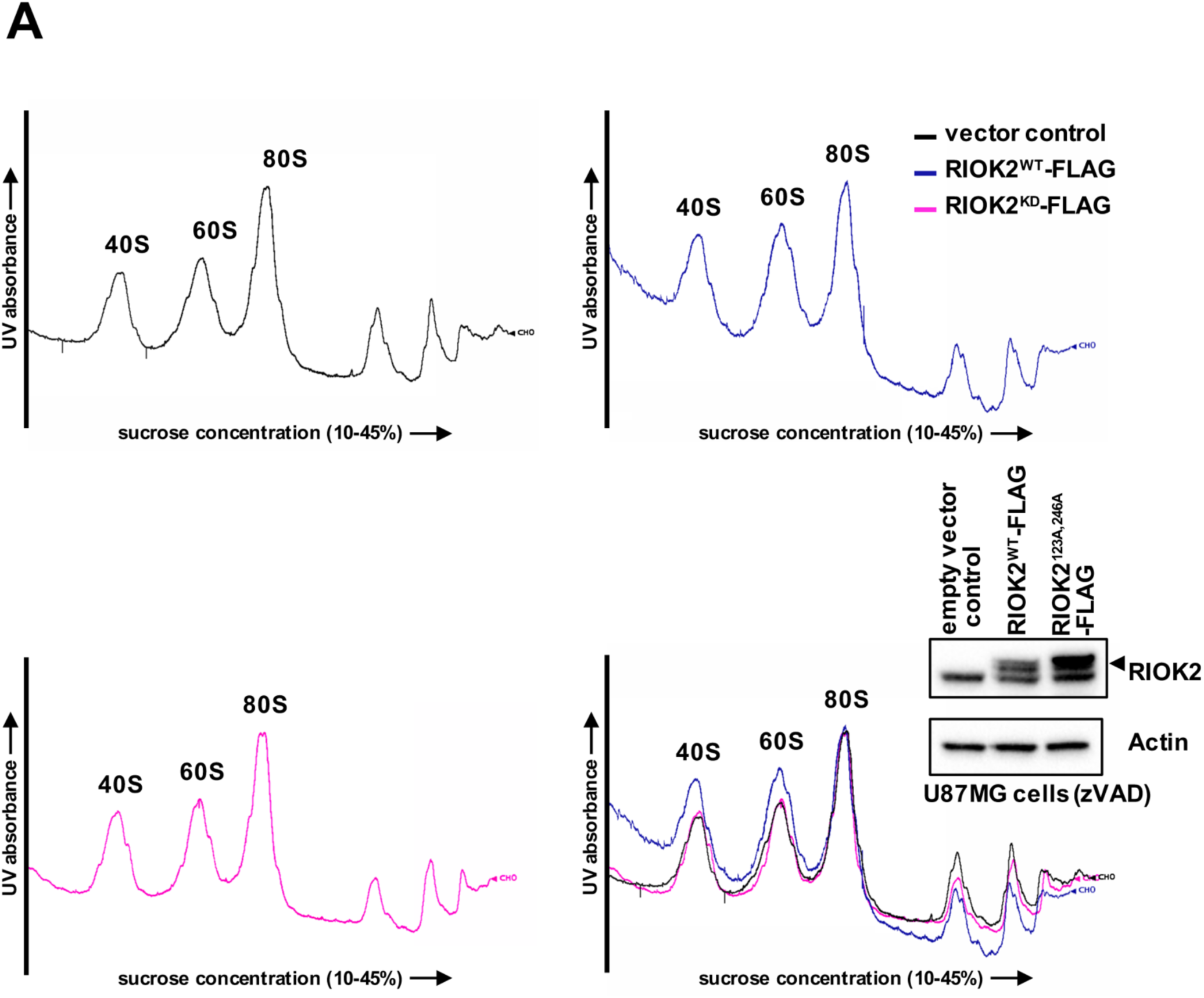
**Overexpressed catalytically inactive RIOK2 does not induce defects in ribosome biogenesis.** A) Analytical sucrose density gradient fractionation of from U87MG GBM cells overexpressing either empty vector control, FLAG-tagged wild-type RIOK2 (RIOK2^WT^-FLAG), or FLAG-tagged catalytically-dead RIOK2 (RIOK2^123A,246A^-FLAG). RIOK2 constructs were induced for 48 hr by doxycycline prior to harvesting cells. RIOK2-FLAG induction efficiency confirmed by immunoblot on total cell lysate (arrow indicates higher molecular weight epitope-tagged RIOK2).10 ODs from each lysate were ultracentrifuged in a sucrose density gradient of 10%-45%, and eluted into several fractions while a UV lamp was used to detect ribosome subunits. Annotated UV traces show that compared to empty vector control U87MG cells (black trace), RIOK2^123A,246A^-FLAG overexpression did not affect levels of 40S and 60S subunits relative to levels of mature 80S monosomes in U87MG cells (magenta trace), whereas RIOK2^WT^-FLAG overexpression slightly increased levels of 40S and 60S subunits relative to levels of mature 80S monosomes in U87MG cells (blue trace). Bottom right shows all UV traces overlaid normalized to 80S peak height.

**Figure S4.**
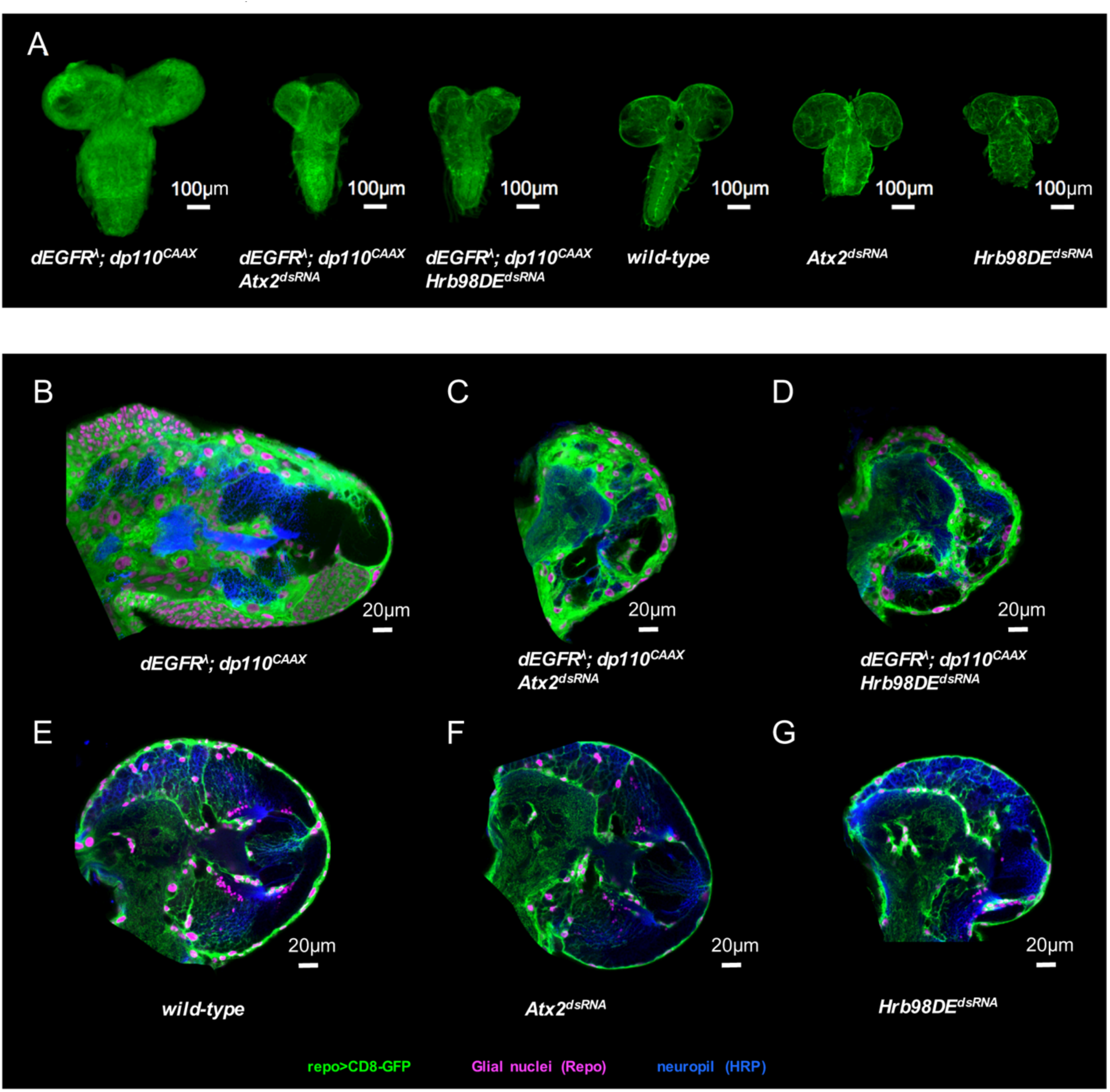
***Drosophila* orthologs of RIOK2 associated RNA-binding proteins are required for EGFR-PI3K glial neoplasia.** (A) Optical projections of whole brain-nerve cord complexes from 3^rd^ instar larvae approximately 6 days old. Dorsal view; anterior up. CD8-GFP (green) labels glial cell bodies. Knockdown of RBPs Atx2 (*repo>dEGFR^λ^;dp110^CAAX^;Atx2^dsRNA^*) or Hrb98DE (*repo>dEGFR^λ^;dp110^CAAX^;Hrb98DE^dsRNA^*) decreased brain overgrowth relative to *repo>dEGFR^λ^;dp110^CAAX^*. Glia-specific knockdown of Atx2 (*repo>Atx2^dsRNA^*) or Hrb98DE (*repo>Hrb98DE^dsRNA^*) induced no obvious defects as compared to wild-type control brains. (B-G) 3 μm optical projections of brain hemispheres, aged-matched 3^rd^ instar larvae. Frontal sections, midway through brains. Anterior up; midline to the left. Repo (magenta) labels glial cell nuclei; CD8-GFP (green) labels glial cell bodies; anti-HRP (blue) counterstains for neurons and neuropil. Knockdown of RBPs (C) Atx2 (*repo>dEGFR^λ^;dp110^CAAX^;Atx2^dsRNA^*) or (D) Hrb98DE (*repo>dEGFR^λ^;dp110^CAAX^;Hrb98DE^dsRNA^*) decreased the numbers of neoplastic glia compared to (B) *repo>dEGFR^λ^;dp110^CAAX^*. Glia-specific knockdown of (F) Atx2 (*repo>Atx2^dsRNA^*) or (G) Hrb98DE (*repo>Hrb98DE^dsRNA^*) induced no drastic differences in glial cell numbers or morphology as compared to (E) wild-type animals.

**Figure S5.**
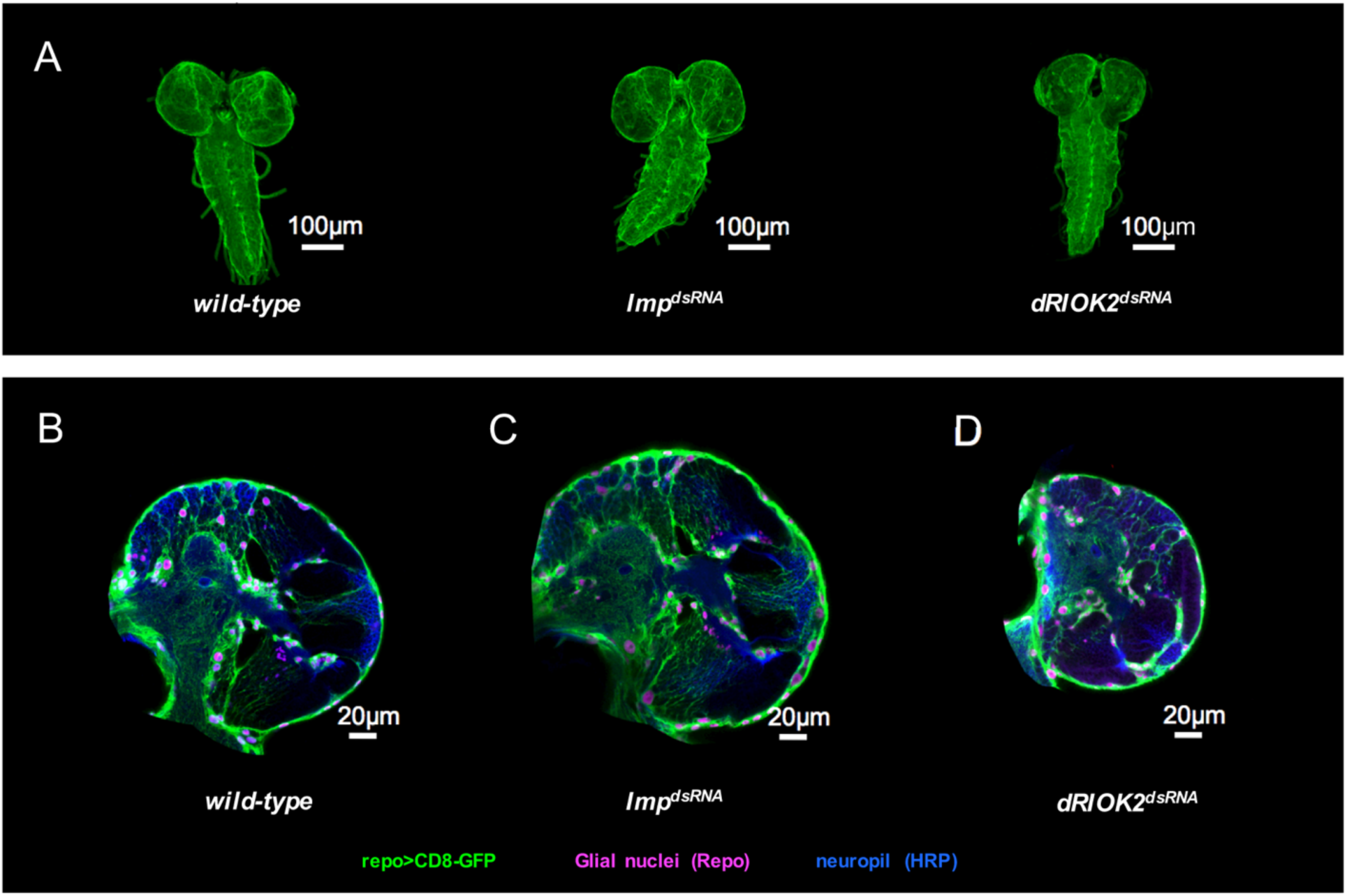
**Imp and dRIOK2 knockdown in wild-type background produced no gross morphological defects in *Drosophila* glia.** (A) Optical projections of whole brain-nerve cord complexes from 3^rd^ instar larvae approximately 6 days old. Dorsal view; anterior up. CD8-GFP (green) labels glial cell bodies. Imp (*repo>Imp^dsRNA^*) or dRIOK2 (*repo>dRIOK2^dsRNA^*) knockdown induced no drastic morphological differences in glia compared to wild-type control animals. (B-D) 3 μm optical projections of brain hemispheres, aged-matched 3^rd^ instar larvae. Frontal sections, midway through brains. Anterior up; midline to the left. Repo (magenta) labels glial cell nuclei; CD8-GFP (green) labels glial cell bodies; anti-HRP (blue) counterstains for neurons and neuropil. Compared to (A) wild-type animals, neither (B) *Imp* knockdown *(repo>Imp^dsRNA^*) nor (D) dRIOK2 knockdown (*repo>dRIOK2^dsRNA^*) showed drastic defects in glial cell numbers (red nuclei) or morphology (green cell bodies).

**Figure S6:**
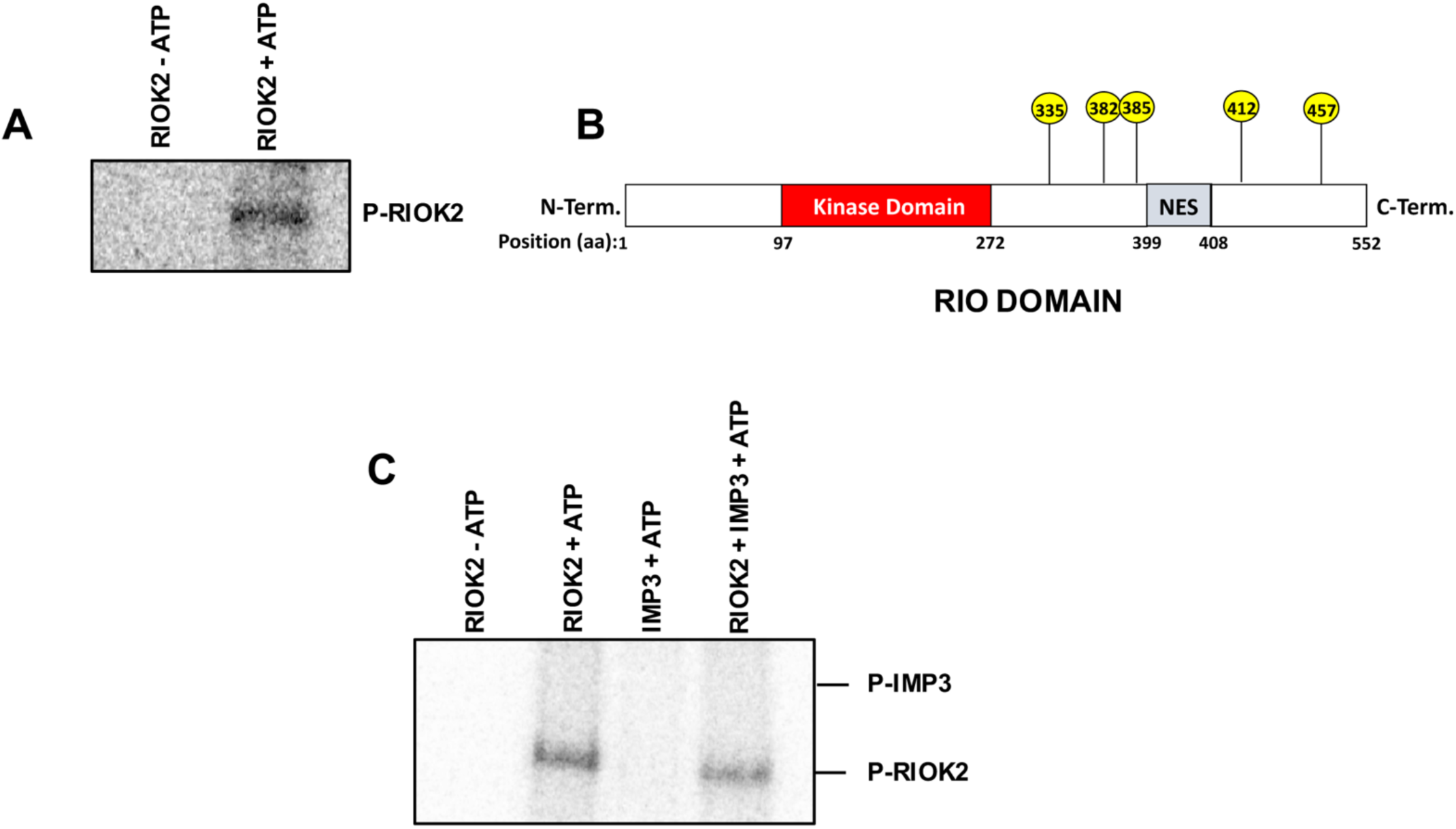
**RIOK2 autophosphorylates itself but does not directly phosphorylate IMP3.** (A) Phospho-image of *in vitro* kinase assay with purified bacterially produced recombinant RIOK2 protein using ^32^P radiolabeled ATP show that RIOK2 is autophosphorylated. (B) Representative image of potential RIOK2 autophosphorylation-sites from an *in vitro* kinase assay using purified bacterially produced labelled with cold ATP, identified using phospho-proteomics. (C) Phospho-image of *in vitro* kinase assay using purified bacterially produced recombinant RIOK2 and/or IMP3 protein mixed together and radiolabeled with ^32^P ATP show that RIOK2 is autophosphorylated, but that RIOK2 does not directly phosphorylate IMP3.

**Figure S7:**
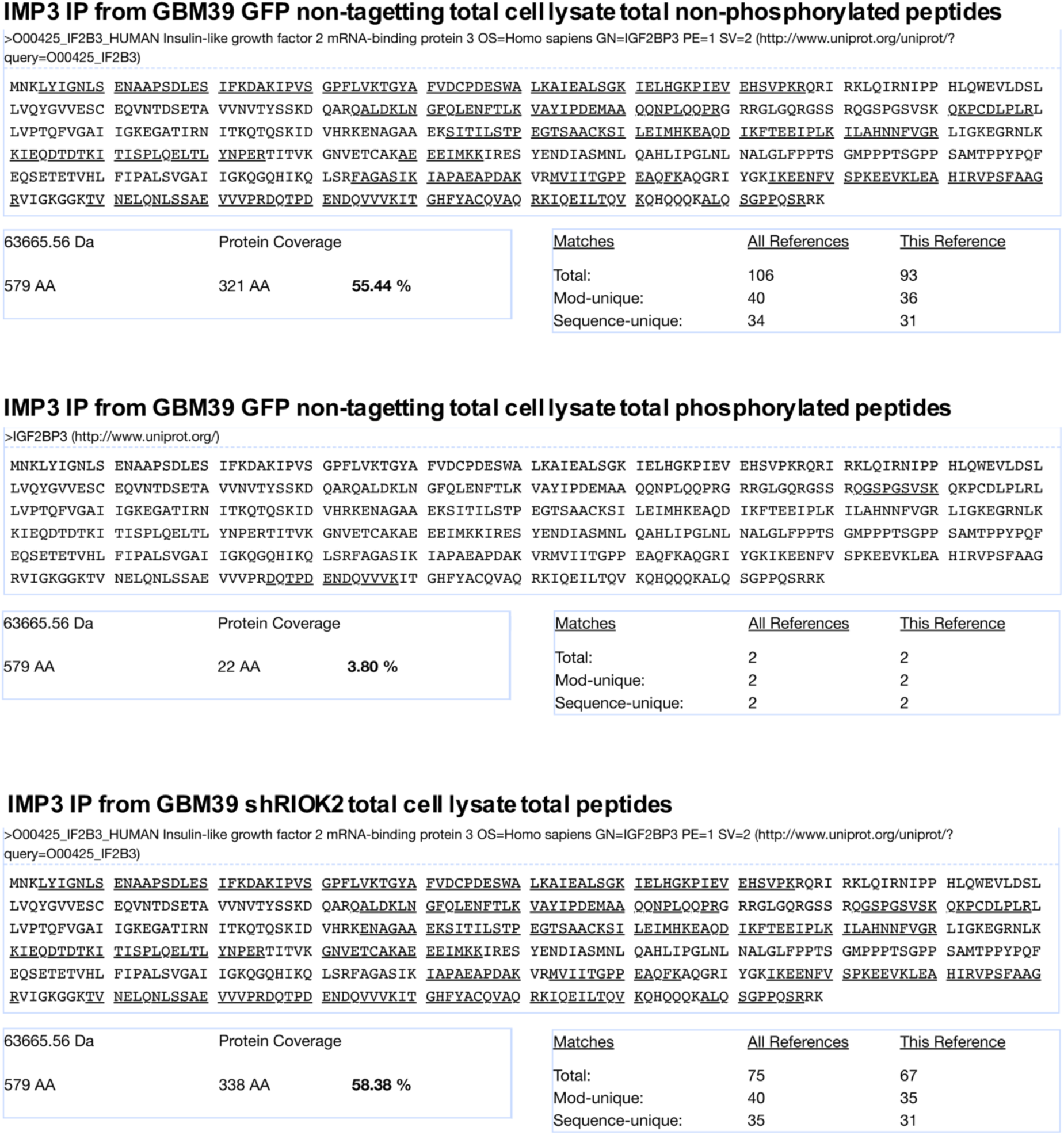
**RIOK2 phospho-proteomic analysis.** Phospho-peptide profiling of endogenous IMP3 upon RIOK2 knockdown. GBM39 cells were treated with verified lentiviral RIOK2 shRNAs. Control cells were infected with a non-targeting GFP control lentivirus. GBM cells were treated with 20 μM zVAD to prevent apoptosis. Cells were harvested 72 hrs after infection, and IMP3 was IPed. RIOK2 knockdown in pre-IP total cell lysates and anti-IMP3 IPs were verified by immunoblots (see Figure 5D). IMP3 was isolated on acrylamide gel slice and profiled for total protein and phosphorylation. The alignment shows total peptide coverage for each IP (residues covered by peptides are underlined in the specific reference alignments shown) and for non-phosphorylated and phosphorylated peptides. No serine/threonine phosphorylated peptides were detected in lysates from cells treated with RIOK2 lentiviral shRNAs.

**Figure S8:**
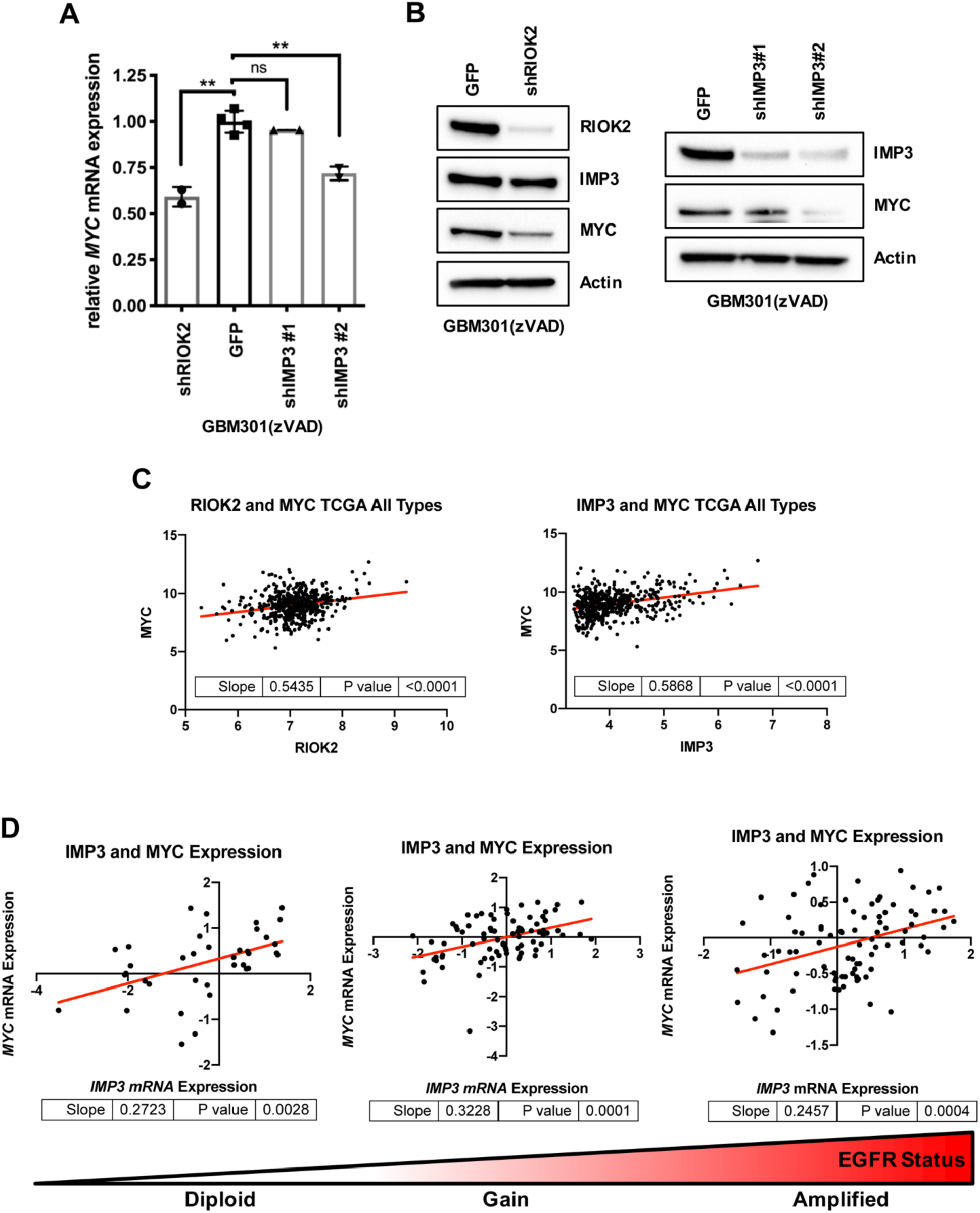
**RIOK2 and IMP3 modulate MYC expression in GBM cells in a conserved pathway.** RIOK2 or IMP3 knockdown was performed in GBM 301cells using verified lentiviral shRNAs. GBM301 cells were treated with 20 μM zVAD to prevent apoptosis. GBM301 cells with RIOK2 and IMP3 knockdown showed reduced expression of endogenous *MYC* mRNA, as measured by qPCR, and MYC protein, as measured by immunoblot, compared to control cells infected with GFP shRNA. The samples shown in this figure were harvested 72 hrs after infection, although by 96 hrs after infection we observed a stronger effect on MYC protein levels as shown in Figure 6C, indicating that MYC mRNA and protein levels drop following a prolonged reduction of IMP3 protein. Two replicates were used per condition. Statistics generated using unpaired two-tailed Student TTESTS, **p<0.01. (C, D) Analysis of TCGA glioma tumor profiling data show significant correlation between increased expression of both *RIOK2* and *IMP3* mRNA and *MYC* mRNA expression across all TCGA glioma grades (C). Increased expression of *IMP3* and *MYC* mRNA are significantly correlated with each other and significantly increased in grade IV gliomas with EGFR gain and amplification compared to tumors diploid for EGFR (D).

**Figure S9:**
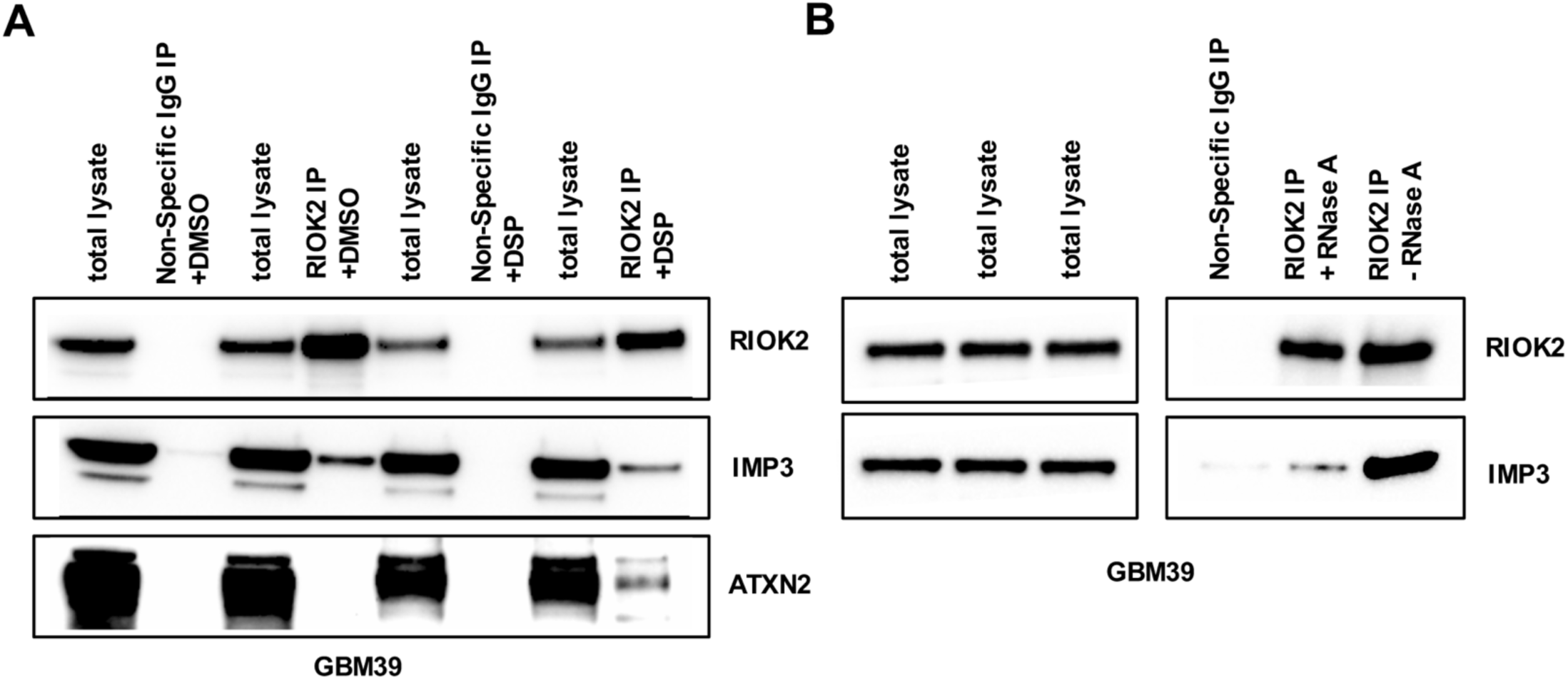
**RIOK2-IMP3 complexes include other RNA-binding proteins and are stabilized by RNA.** A). GBM39 treated with either DMSO or DSP crosslinker (1 mM) for 2 hrs prior to IPs with either a RIOK2 antibody or a non-specific IgG control. Immunoblots show enrichment of endogenous ATXN2 protein in the RIOK2 IP samples treated with DSP. B). GBM39 total cell lysates pre-treated with RNAse A (100µg/ml) prior to IPs with either a RIOK2 antibody or a non-specific IgG control. Immunoblots show decreased levels of IMP3 protein co-IPed with RIOK2 from the lysate treated with RNase A.

**Table S1:** **Proteomic approach to identify endogenous RIOK2 binding proteins.** Proteomic analysis of IPs from U87MG-EGFR^vIII^ cells and GBM301 cells performed with antibodies against endogenous RIOK2 or a non-specific IgG control. Values are reported in spectral counts (PSM) or number of unique peptides. Spectral counts are the total number of spectra identified for a protein and is a relative measurement of the quantity of a given protein, while the number of unique peptides refers to the number of peptides that matches the protein in question. Names of the human genes are given along with the name of the associated fly homolog.

| GENE | FLY Homolog | U87MG-EGFR<sup>viii</sup> |  |  |  | GBM301 |  |  |  |
| --- | --- | --- | --- | --- | --- | --- | --- | --- | --- |
|  |  | Non-Specific IgG IP |  | RIOK2 IP |  | Non-Specific IgG IP |  | RIOK2 IP |  |
|  |  | Spectral Counts | Unique Peptide | Spectral Counts | Unique Peptide | Spectral Counts | Unique Peptide | Spectral Counts | Unique Peptide |
| *LTV1* | *CG7686* | 0 | 0 | 21 | 13 | 0 | 0 | 23 | 14 |
| *NOB1* | *CG2972* | 0 | 0 | 12 | 9 | 2 | 2 | 10 | 8 |
| *PNO1* | *lethal (1) G0004* | 0 | 0 | 18 | 7 | 2 | 2 | 18 | 10 |
| *G3BP2* | *rasputin* | 0 | 0 | 8 | 6 | 4 | 2 | 13 | 8 |
| *ILF3* | *Zn72D* | 0 | 0 | 13 | 11 | 17 | 12 | 29 | 22 |
| *RIOK2* | *RIOK2* | 0 | 0 | 22 | 11 | 0 | 0 | 14 | 12 |
| *IMP3* | *Imp* | 0 | 0 | 8 | 6 | 0 | 0 | 5 | 3 |
| *BYSL* | *bys* | 0 | 0 | 17 | 12 | 2 | 2 | 18 | 11 |
| *TSR1* | *CG7338* | 0 | 0 | 30 | 22 | 0 | 0 | 28 | 19 |
| *SEC23IP* | *PAPLA1* | 1 | 1 | 17 | 12 | 1 | 1 | 5 | 5 |
| *GIGYF2* | *CG11148* | 1 | 1 | 9 | 5 | 1 | 1 | 14 | 12 |
| *G3BP1* | *rasputin* | 4 | 4 | 27 | 13 | 7 | 7 | 3 | 1 |
| *PABPC4* | *pAbp* | 6 | 1 | 28 | 8 | 10 | 8 | 17 | 6 |
| *ATXN2L* | *Atx2* | 2 | 1 | 8 | 6 | 4 | 3 | 10 | 8 |
| *RRBP1* | *alt* | 5 | 4 | 19 | 1 | 28 | 28 | 57 | 47 |
| *DDX1* | *Ddx1* | 8 | 6 | 26 | 12 | 7 | 6 | 21 | 15 |
| *HNRNPUL1* | *CG30122* | 1 | 1 | 3 | 1 | 13 | 11 | 6 | 4 |
| *DDX5* | *CG10077* | 10 | 4 | 29 | 13 | 7 | 3 | 19 | 9 |
| *HNRNPA3* | *Hrb98DE, Hrb87F* | 5 | 4 | 13 | 1 | 11 | 8 | 13 | 11 |
| *DDX17* | *CG10077* | 13 | 7 | 19 | 8 | 7 | 3 | 21 | 12 |
| *HNRNPL* | *smooth* | 4 | 3 | 5 | 5 | 2 | 2 | 6 | 6 |

**Table S2.** **IMP3 shows reduced binding with catalytically inactive RIOK2.** Proteomic analysis of RIOK2^WT^-FLAG or catalytically-dead RIOK2^123A,246A^-FLAG immunoprecipitates from U87MG cells, both with an N-terminal FLAG epitope tag, and empty vector control cells. RIOK2-FLAG constructs were overexpressed in U87MG cells using doxycycline inducible vectors. Cells were treated with 8 μg/ml of doxycycline and incubated for 72 hrs and then lysed under hypotonic conditions. Using M2 antibodies to the FLAG epitope tag, IPs were performed on total cell lysates from all three samples, and subjected to proteomic analysis. Values are reported in spectral counts (PSM) and number of unique peptides. Spectral counts are the total number of spectra identified for a protein and is a relative measurement of the quantity of a given protein, while the number of unique peptides refers to the number of peptides that matches the protein in question.

| Table S2: Proteomics approach to identify RIOK2-FLAG binding proteins |  |  |  |  |  |  |
| --- | --- | --- | --- | --- | --- | --- |
| Human Gene | Vector Control |  | RIOK2<sup>WT</sup> FLAG IP |  | RIOK2<sup>KD</sup> FLAG IP |  |
|  | Spectral Counts | Unique Peptide | Spectral Counts | Unique Peptide | Spectral Counts | Unique Peptide |
| *LTV1* | 0 | 0 | 8 | 6 | 11 | 7 |
| *NOB1* | 0 | 0 | 7 | 4 | 8 | 7 |
| *PNO1* | 1 | 1 | 11 | 5 | 12 | 7 |
| *G3BP2* | 4 | 3 | 6 | 4 | 6 | 5 |
| *ILF3* | 4 | 4 | 5 | 3 | 11 | 5 |
| ***RIOK2*** | 0 | 0 | 78 | 23 | 78 | 23 |
| ***IMP3*** | 2 | 2 | 5 | 2 | 2 | 1 |
| *BYSL* | 0 | 0 | 4 | 3 | 2 | 2 |
| *TSR1* | 1 | 1 | 18 | 11 | 6 | 6 |
| *SEC23IP* | 3 | 3 | 5 | 5 | 8 | 5 |
| *GIGYF2* | 2 | 2 | 3 | 3 | 2 | 2 |
| *G3BP1* | 11 | 7 | 14 | 8 | 17 | 9 |
| *PABPC4* | 9 | 2 | 11 | 1 | 15 | 4 |
| *ATXN2L* | 5 | 4 | 7 | 5 | 10 | 9 |
| *RRBP1* | 8 | 7 | 13 | 1 | 14 | 1 |
| *DDX1* | 8 | 6 | 11 | 5 | 15 | 9 |
| *HNRNPUL1* | 1 | 1 | 5 | 3 | 4 | 4 |
| *DDX5* | 23 | 14 | 38 | 17 | 29 | 16 |
| *HNRNPA3* | 13 | 9 | 17 | 1 | 7 | 6 |
| *DDX17* | 9 | 6 | 21 | 9 | 17 | 7 |
| *HNRNPL* | 9 | 8 | 11 | 8 | 17 | 11 |

**Table S3.**
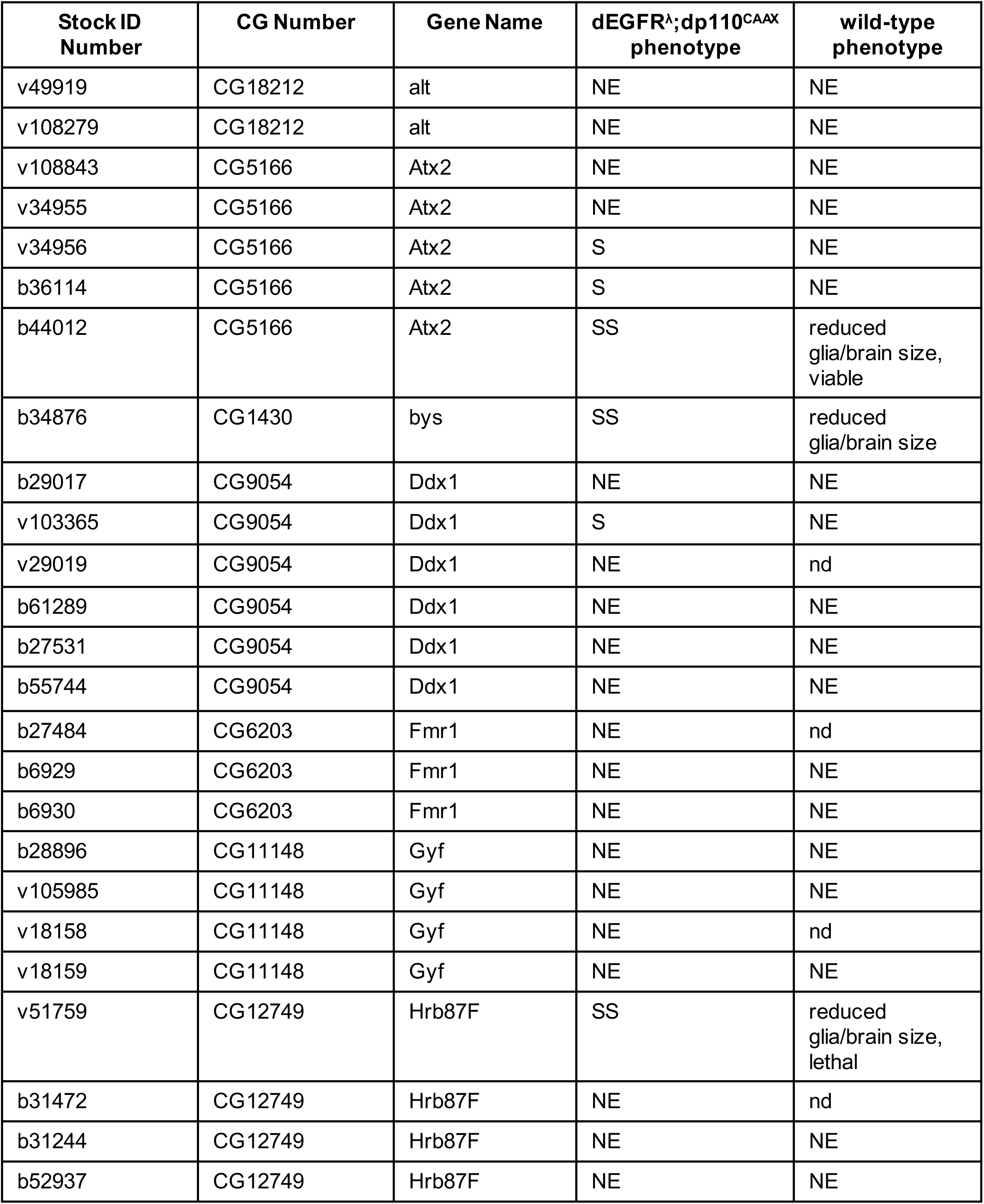
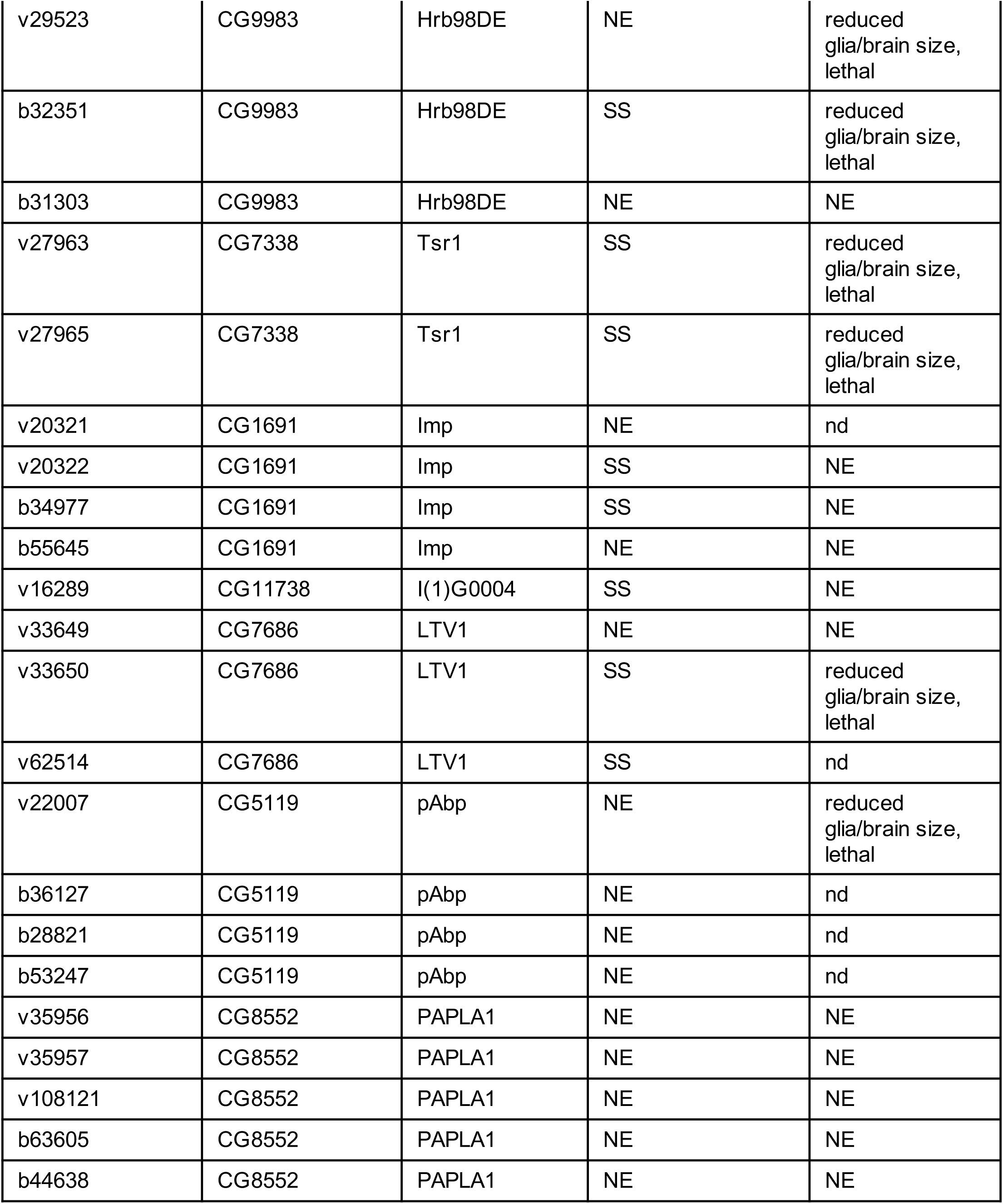
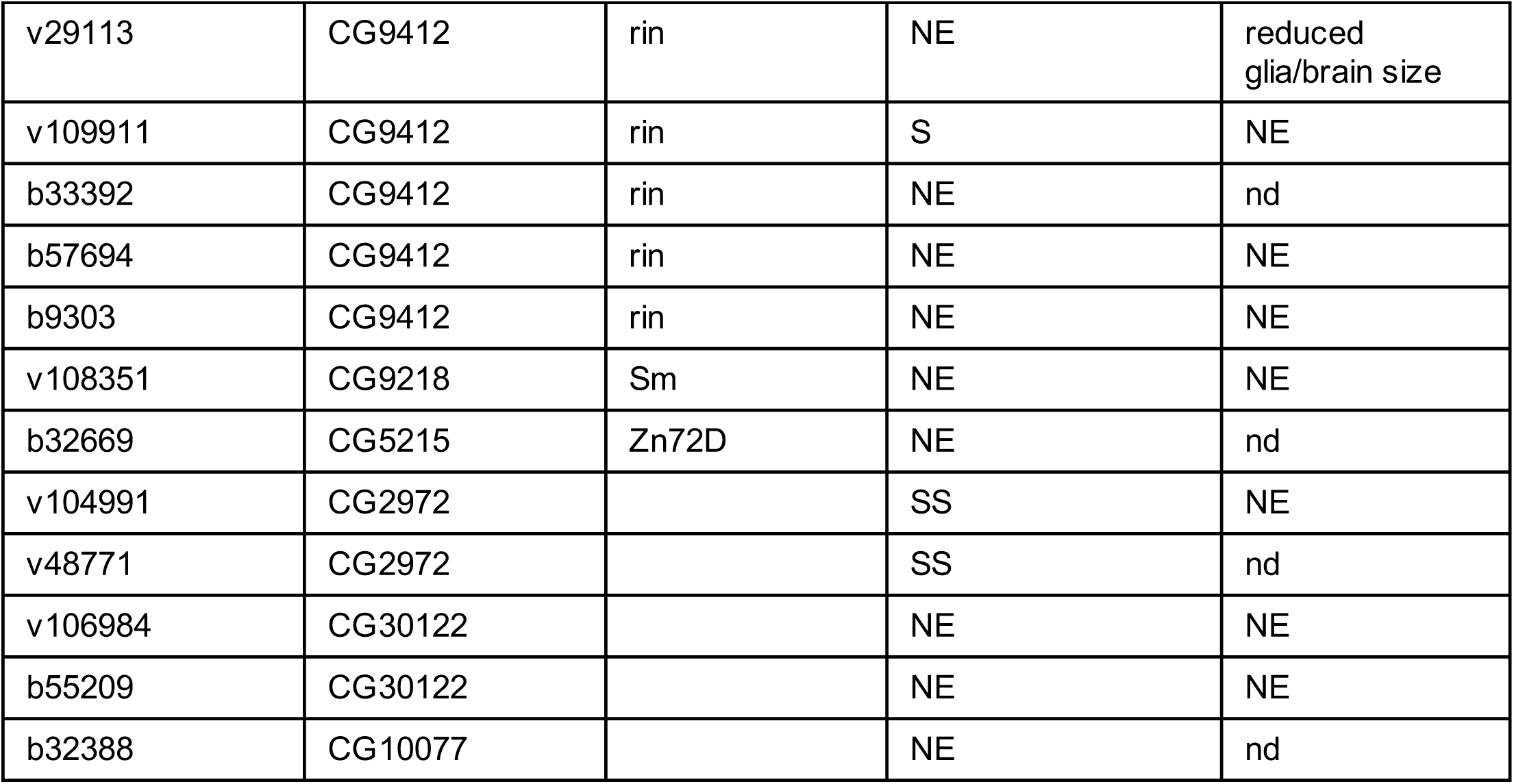
**RNAi screen of *Drosophila* orthologs for RIOK2 binding proteins.** UAS-dsRNA stocks were obtained for *Drosophila* orthologs of RIOK2 binding proteins and tested to determine the effects of RNAi knockdown of their target genes on neoplastic glia using the *repo>dEGFR^λ^;dp110^CAAX^* GBM model and, for comparison, on wild-type glia using *repo-Gal4*. Not all UAS-dsRNA stocks tested have been validated for their RNAi efficacy. VDRC stock ID numbers prefaced with a “v,” Bloomington stock ID numbers are prefaced by a “b”. Key to genetics interactions: WS: weak suppressor, S: moderate suppressor, SS: strong suppressor; NE: no effect indicates that there were no obvious phenotypic differences between dsRNA and control animals; nd indicates no determined.

| **Stock ID Number** | **CG Number** | **Gene Name** | **dEGFR<sup>Δ</sup>;dp110<sup>CAAX</sup> phenotype** | **wild-type phenotype** |
| --- | --- | --- | --- | --- |
| v49919 | CG18212 | alt | NE | NE |
| v108279 | CG18212 | alt | NE | NE |
| v108843 | CG5166 | Atx2 | NE | NE |
| v34955 | CG5166 | Atx2 | NE | NE |
| v34956 | CG5166 | Atx2 | S | NE |
| b36114 | CG5166 | Atx2 | S | NE |
| b44012 | CG5166 | Atx2 | SS | reduced glia/brain size, viable |
| b34876 | CG1430 | bys | SS | reduced glia/brain size |
| b29017 | CG9054 | Ddx1 | NE | NE |
| v103365 | CG9054 | Ddx1 | S | NE |
| v29019 | CG9054 | Ddx1 | NE | nd |
| b61289 | CG9054 | Ddx1 | NE | NE |
| b27531 | CG9054 | Ddx1 | NE | NE |
| b55744 | CG9054 | Ddx1 | NE | NE |
| b27484 | CG6203 | Fmr1 | NE | nd |
| b6929 | CG6203 | Fmr1 | NE | NE |
| b6930 | CG6203 | Fmr1 | NE | NE |
| b28896 | CG11148 | Gyf | NE | NE |
| v105985 | CG11148 | Gyf | NE | NE |
| v18158 | CG11148 | Gyf | NE | nd |
| v18159 | CG11148 | Gyf | NE | NE |
| v51759 | CG12749 | Hrb87F | SS | reduced glia/brain size, lethal |
| b31472 | CG12749 | Hrb87F | NE | nd |
| b31244 | CG12749 | Hrb87F | NE | NE |
| b52937 | CG12749 | Hrb87F | NE | NE |
| v29523 | CG9983 | Hrb98DE | NE | reduced glia/brain size, lethal |
| b32351 | CG9983 | Hrb98DE | SS | reduced glia/brain size, lethal |
| b31303 | CG9983 | Hrb98DE | NE | NE |
| v27963 | CG7338 | Tsr1 | SS | reduced glia/brain size, lethal |
| v27965 | CG7338 | Tsr1 | SS | reduced glia/brain size, lethal |
| v20321 | CG1691 | Imp | NE | nd |
| v20322 | CG1691 | Imp | SS | NE |
| b34977 | CG1691 | Imp | SS | NE |
| b55645 | CG1691 | Imp | NE | NE |
| v16289 | CG11738 | l(1)G0004 | SS | NE |
| v33649 | CG7686 | LTV1 | NE | NE |
| v33650 | CG7686 | LTV1 | SS | reduced glia/brain size, lethal |
| v62514 | CG7686 | LTV1 | SS | nd |
| v22007 | CG5119 | pAbp | NE | reduced glia/brain size, lethal |
| b36127 | CG5119 | pAbp | NE | nd |
| b28821 | CG5119 | pAbp | NE | nd |
| b53247 | CG5119 | pAbp | NE | nd |
| v35956 | CG8552 | PAPLA1 | NE | NE |
| v35957 | CG8552 | PAPLA1 | NE | NE |
| v108121 | CG8552 | PAPLA1 | NE | NE |
| b63605 | CG8552 | PAPLA1 | NE | NE |
| b44638 | CG8552 | PAPLA1 | NE | NE |

| Table S3: RNAi screen of *Drosophila* orthologs for RIOK2 binding proteins |  |  |  |  |
| --- | --- | --- | --- | --- |
| Stock ID Number | CG Number | Gene Name | dEGFR<sup>Δ</sup>;dp110<sup>CAAX</sup> Phenotype | Wild-Type Phenotype |
| v29113 | CG9412 | rin | NE | reduced glia/brain size |
| v109911 | CG9412 | rin | S | NE |
| b33392 | CG9412 | rin | NE | nd |
| b57694 | CG9412 | rin | NE | NE |
| b9303 | CG9412 | rin | NE | NE |
| v108351 | CG9218 | Sm | NE | NE |
| b32669 | CG5215 | Zn72D | NE | nd |
| v104991 | CG2972 |  | SS | NE |
| v48771 | CG2972 |  | SS | nd |
| v106984 | CG30122 |  | NE | NE |
| b55209 | CG30122 |  | NE | NE |
| b32388 | CG10077 |  | NE | nd |

